# A Preoptic Neurocircuit That Modulates Metabolic Flexibility

**DOI:** 10.64898/2026.01.15.699760

**Authors:** Julian M. Roessler, Matthew Alkire, Nathan Nigrin, Haorui Wang, Christopher M. Reid, Marissa D. Cortopassi, Millenia Waite, Brooke Linnehan, Eric C. Griffith, Mollie Madigan, Tenzin Kunchok, Alexander S. Banks, Fabian Schulte, Bukyung Kim, Bo-Yeon Kim, Jason K. Kim, Siniša Hrvatin

## Abstract

Precise, dynamic control of metabolic fuel usage in response to environmental challenges such as altered food availability or temperature change is essential for animal survival. In mammals, metabolic flexibility—the capacity to shift cellular metabolism between carbohydrate and fatty acid oxidation—is understood to be largely regulated by circulating hormones such as insulin and glucagon. However, the role of the central nervous system in coordinating fuel selection and tissue metabolic tuning remains underexplored. Here, we investigated the mechanisms that mediate metabolic reprogramming following the acute activation of torpor-associated glutamatergic Adcyap1+ torpor-regulating neurons in the anteroventral preoptic area (avPOA^Vglut2/PACAP^). The activation of these neurons rapidly shifts whole-body fuel use from glucose to fatty acids, irrespective of fuel/food availability. This shift is associated with reduced glucose utilization stemming from the transient induction of selective insulin resistance in skeletal muscle. We find that this reduction in skeletal muscle glucose metabolism does not require direct muscle innervation but is rather mediated in part via corticosterone. In contrast to their activation, avPOA^Vglut2/PACAP^ neuronal silencing results in improved glucose tolerance, demonstrating powerful bidirectional control of tissue-specific glucose metabolism, whole-body glucose levels, and fuel usage. Together, our findings uncover a novel POA -skeletal muscle pathway that dynamically controls glucose utilization and metabolic flexibility.

## Introduction

Dynamic control of metabolism is essential for homeostatic maintenance in response to changing environments, including altered temperature, food availability, and stress. For example, in the context of nutrient deficit or increased energetic demands, fuel usage shifts from mixed fatty acid and carbohydrate oxidation to near complete reliance on fat metabolism to preserve glucose for use by the brain and prevent hypoglycemia^1,2^. Metabolic inflexibility, the inability to efficiently switch between these fuel sources, has been linked to obesity, diabetes, and metabolic disease^3,4^.

Baseline regulation of blood sugar and fuel usage is largely controlled by the circulating pancreatic hormones insulin and glucagon, with insulin resistance representing a key aspect of disease-associated metabolic inflexibility^3–5^. While these hormonal regulators of metabolic flexibility and their peripheral effectors have been extensively studied, less is known concerning the role of the central nervous system (CNS) in dynamic top-down metabolic control. The ventromedial hypothalamus (VMH) is the primary nucleus in the brain known to control systemic glucose usage. Activation of counterregulatory steroidogenic factor 1 (SF-1)- or Cholecystokinin B receptor (CCKBR)-expressing neurons in the VMH drives increased levels of systemic glucose and peripheral tissue insulin sensitivity^6,7^. In addition, brain-body signaling via the hypothalamic-pituitary-adrenal (HPA) axis has long been known to play a crucial role in stress-induced metabolic responses, influencing insulin sensitivity and energy balance^8,9^. Yet, which neuronal populations and brain-body axes mediate dynamic metabolic reprogramming in response to various environmental challenges remain to be fully characterized.

In response to prolonged periods of food deprivation, a wide range of mammalian species, including laboratory mice (*Mus musculus*), have evolved the capacity to suspend or override canonical fasting metabolism to enter protective hypometabolic states such as daily torpor and hibernation^10^. Fasting-induced daily torpor in mice is characterized by a reduction of metabolic rate by approximately 60%^2,10^ and an accompanying reduction in body temperature from 37°C to near ambient levels. It has long been appreciated that entry into and maintenance of these states requires utilization of internal fat stores as a primary energy source, shifting metabolism almost entirely towards fatty acid oxidation^2^. Recent work from our group and others has shown that torpor entry is centrally controlled by the activity of glutamatergic neurons within the anteroventral preoptic area of the hypothalamus (avPOA^Vglut2^ neurons). The activation of this neuronal population results in a dramatic reduction in energy expenditure even in the absence of caloric restriction, suggesting that these neurons are powerful modulators of systemic metabolic flexibility ^11,12,13^. Discrete activation of this circuit thus provides a means to dissect a central pathway for controlling metabolic flexibility, without the confounding metabolic effects of prolonged fasting.

Here, we leverage the power of targeted opto- and chemogenetic circuit manipulation to probe the metabolic effects of avPOA^Vglut2^ neuronal activation in the mouse and trace downstream consequences for peripheral organ-specific glucose regulation. We show that avPOA^Vglut2^ neuronal activation drives a preferential reduction in whole-body glucose metabolism and a shift to a fatty acid oxidative state stemming from acute tissue-specific skeletal muscle and brown adipose tissue insulin resistance. Moreover, acute inhibition of this population improves glucose tolerance, nominating avPOA^Vglut2^ neurons as powerful bidirectional regulators of glucose homeostasis. The resulting reduction in skeletal muscle glucose uptake and systemic glucose intolerance is not dependent on the decrease in core body temperature, nor direct muscle innervation, but rather is hormonally driven by the adrenal gland secretion of corticosterone. Furthermore, these effects are recapitulated by the Adcyap1^+^ (PACAP-expressing) subpopulation of avPOA^Vglut2^ neurons. Together, these findings identify avPOA^Vglut2^ neurons as a significant unappreciated CNS glucoregulatory population intimately involved in the regulation of the overall metabolic state. Targeted manipulation of this neuronal population and its downstream effectors may thus prove relevant to the amelioration of disorders of metabolic inflexibility, such as obesity, diabetes, and metabolic syndrome.

## Results

### avPOA^Vglut2^ neurons control systemic metabolic fuel usage independent of changes in body temperature

Torpor is canonically understood as a strategy for energetic conservation during times of nutrient deprivation^2^. While it is well-appreciated that fasting induces a shift from carbohydrate oxidation (CHO) to fatty acid oxidation (FAO)^14^, how torpor entry affects CHO versus FAO utilization relative to surrounding periods of fasting at euthermic body temperatures remains less well characterized. To monitor shifting fuel usage during fasting-induced torpor, 8-week-old wild-type C57Bl/6J mice were implanted intraperitoneally with telemetric temperature probes, fasted in metabolic cages for 24 hours, and monitored for changes in body temperature (T_b_), gas exchange, and respiratory exchange ratio (RER) — an established measure of CHO versus FAO utilization (**Figure 1a)**. RER levels were found to decrease substantially during periods of torpor relative to the fed state and surrounding periods of euthermic fasting (**Figure 1a, b)**, consistent with an acute shift away from CHO and towards FAO during torpor.

**Figure 1:**
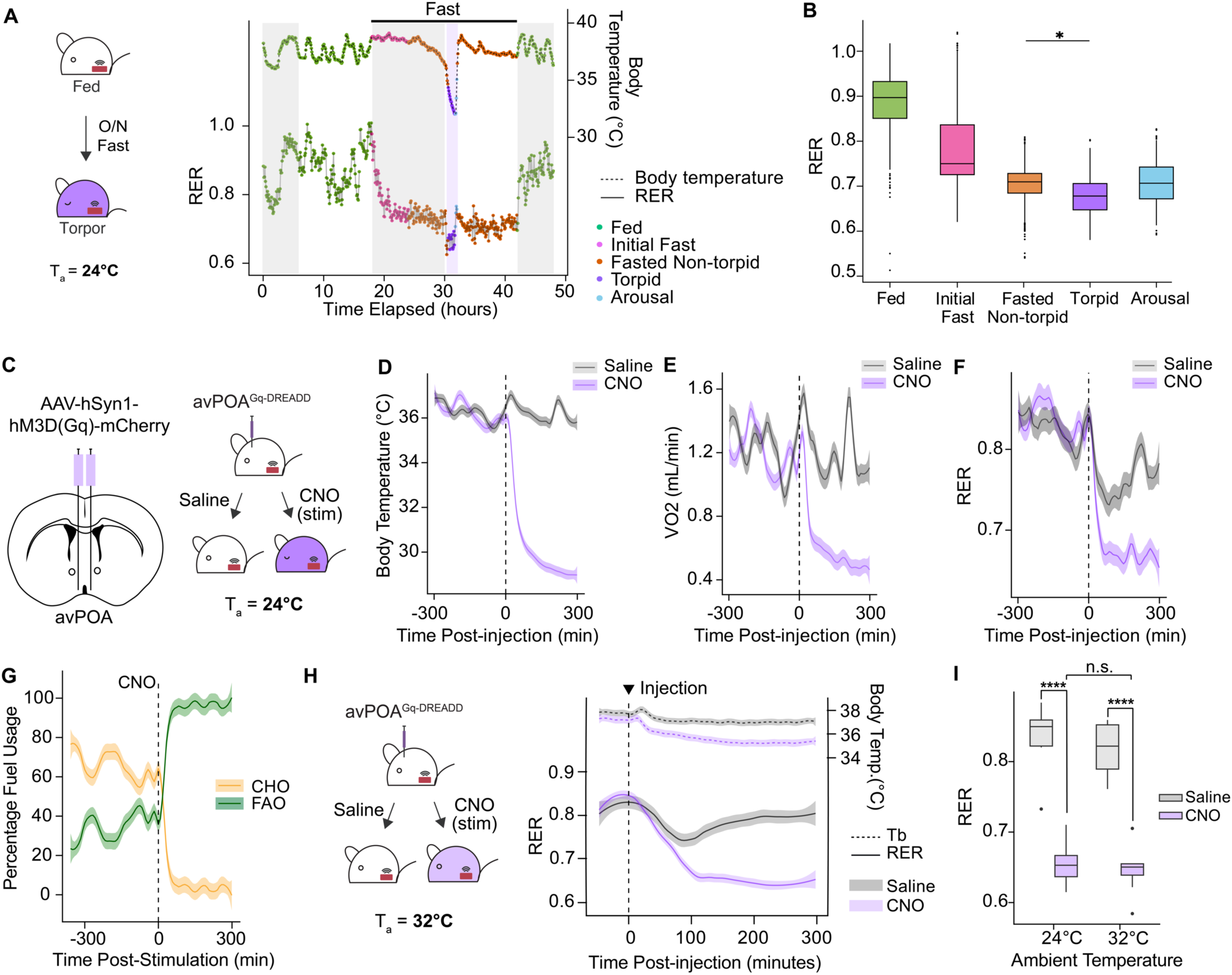
avPOA^Vglut2^ neurons regulate fuel usage in natural and induced torpor. **(A)** *(Left)* Schematic for respirometry experiments involving natural torpor. Probe-implanted wild-type C57Bl6/J mice were monitored throughout a 24-hour fast (ZT12-ZT12, denoted by horizontal line). (*Right)* Representative traces of respiratory exchange ratio (RER, bottom, solid line) and body temperature (T_b_, top, dashed line) for an animal undergoing natural fasting-induced torpor. Traces were classified into distinct states based on food availability and T_b_: *Fed*, ad libitum access to chow; *Initial Fast*, <6 h post-food withdrawal; *Fasted Non-torpid*, >6 h fasting and T_b_ > 34.5°C; Torpid –fasted, T_b_ < 34.5°C, and ΔT_b_ < 0; *Arousal*, T_b_ < 35.5°C and ΔT_b_ > 0. Shaded gray portions denote dark cycle times (ZT12-24). **(B)** State-dependent quantification of RER for fasted mice as in (A) (n = 8 animals, 4 male, 4 female, * = p < 0.05). **(C)** (*Left)* Schematic for bilateral stereotactic transduction of avPOA neurons with Gq-DREADD. *(Right)* Experimental paradigm for stimulating avPOA-transduced animals with either saline or CNO (1 mg/kg) at T_a_ = 24°C. (**D-F)** Quantification of T_b_ (**D**), VO_2_ (**E**), and RER (**F**) in Gq-DREADD-expressing mice for 300 minutes pre- and post-CNO stimulation (dashed vertical line). Traces and shading indicate mean ± SEM (n = 8 animals). **(G)** Calculation of carbohydrate oxidation (CHO) and fatty acid oxidation (FAO) percentages for 300 minutes pre- and post-CNO injection (dashed vertical line) based on the method of Frayn (see Methods). **(H)** (*Left*) Following acclimation at elevated T_a_ (32°C), Gq-DREADD-expressing mice were administered either saline or CNO and monitored for gas exchange and T_b_. (*Right)* RER and T_b_ changes in Gq-DREADD-expressing mice in response to CNO stimulation or saline control. Dashed vertical line indicates timing of the injection. Ambient temperature denoted by horizontal dashed line (T_a_ = 32°C). **(I)** Quantitation of RER 300 min following CNO or saline control injections at T_a_ = 24°C or 32°C (**** = p < 0.00005, n = 8 mice).

We previously demonstrated that chemogenetic stimulation of avPOA neurons phenocopies the decreased T_b_ and metabolic rate observed during natural torpor, thereby providing a model to decouple avPOA neuronal activity from fasting-induced metabolic changes^11^. To determine the metabolic consequences of avPOA neuronal activation, we stereotactically introduced a neuronally targeted adeno-associated virus (AAV) expressing Gq-DREADD^15^ (AAV-hSyn-hM3D(Gq)-mCherry) into the avPOA of C57Bl/6J mice (**Figure 1c**). The Gq-DREADD-activating ligand Clozapine-N-Oxide (CNO) was administered intraperitoneally to these non-fasted animals, resulting in substantial decreases in both body temperature and metabolic rate following CNO injection as compared to saline-injected control animals (**Figures 1d-e, Supplementary Figures 1a,b**). These decreases were accompanied by dramatic reductions in other metabolic parameters, including food intake, VCO_2_, and VO_2_ (**Supplementary Figures 1b-f**), as well as a marked shift in RER, from 0.86±0.01 to 0.66±0.02 (**Figure 1d**), consistent with a shift in metabolic fuel usage from mixed metabolism (carbohydrate and fat) to primarily fat catabolism^3,16^. Moreover, utilizing the method of Frayn^16^ to quantify the relative contribution of CHO and FAO to overall energy expenditure, we confirmed that upon avPOA stimulation animals transition from a mixed fuel state to nearly exclusively using FAO for energy generation (**Figure 1g**). Together, these results demonstrate that avPOA^Vglut2^ neuronal activation alone is sufficient to induce the alterations in fuel usage, independent of caloric deprivation.

To determine whether the observed Gq-DREADD-driven RER effect was mediated by decreased T_b_, we increased the ambient temperature (T_a_) from 24°C to 32°C, reasoning that increased T_a_ would prevent the T_b_ decrease accompanying avPOA^Vglut2^ neuron activation. Indeed, we observed that the average T_b_ of CNO-stimulated animals at this elevated ambient temperature was 35.7±0.24°C as compared with 29.8±0.22°C when stimulated at T_a_ = 24°C (**Figure 1h, Supplementary Figure 1e**). Strikingly, despite the absence of a large T_b_ drop, stimulated animals exhibited a dramatic decrease in RER nearly identical to that observed at 24°C (**Figure 1h**), indicating that the observed metabolic shift is not a secondary consequence of T_b_ reduction.

### avPOA^Vglut2^ neurons regulate blood glucose homeostasis and drive metabolic inflexibility

To gain deeper insight into the metabolic consequences of avPOA neuronal stimulation, we performed polar serum metabolomics analyses to identify changes in circulating metabolites accompanying avPOA stimulation. Having previously identified glutamatergic avPOA (avPOA^Vglut2^) neurons as the primary drivers of torpid hypometabolism, we sought to target this neuronal population more specifically. To this end, we transduced the avPOA of telemetric probe-implanted *Vglut2-IRES-Cre* animals with AAV expressing either a Cre-dependent Gq-DREADD or mCherry control. After collecting tail blood to establish a baseline for comparison, animals were administered CNO, and blood samples were collected 15, 30, 60, and 120 minutes post-injection, while continuously measuring T_b._ We then performed plasma separation and polar metabolome analysis on these samples, examining longitudinal changes in the abundance of polar plasma metabolites in Gq-DREADD mice as compared with their mCherry counterparts (**Figure 2a**). This analysis identified 4 significantly increased and 34 significantly decreased polar plasma metabolites across all time points in avPOA^Vglut2^ neuron-stimulated animals (**Figure 2b**). Observed changes involved significant decreases in key glycolytic and citric acid cycle metabolites, including lactate, pyruvate, and 3-phosphoglycerate, accompanied by increases in beta-hydroxybutyrate (**Figure 2b, Supplementary Figures 2a-d**). These changes were monotonic across the duration of the experiment, suggesting continued suppression of glucose metabolism (**Supplementary Figure 2a**), and were consistent with a state in which low blood glucose levels lead to decreased glycolysis and a compensatory increase in FAO. However, instead of a decrease in blood glucose levels, we observed an increase in serum glucose (**Figure 2a**), suggesting that avPOA^Vglut2^ stimulation suppresses glucose metabolism downstream of glucose availability. This suppression of glucose metabolism generates a metabolically inflexible state, wherein animals rely nearly completely on fatty acid oxidation for energy production.

**Figure 2:**
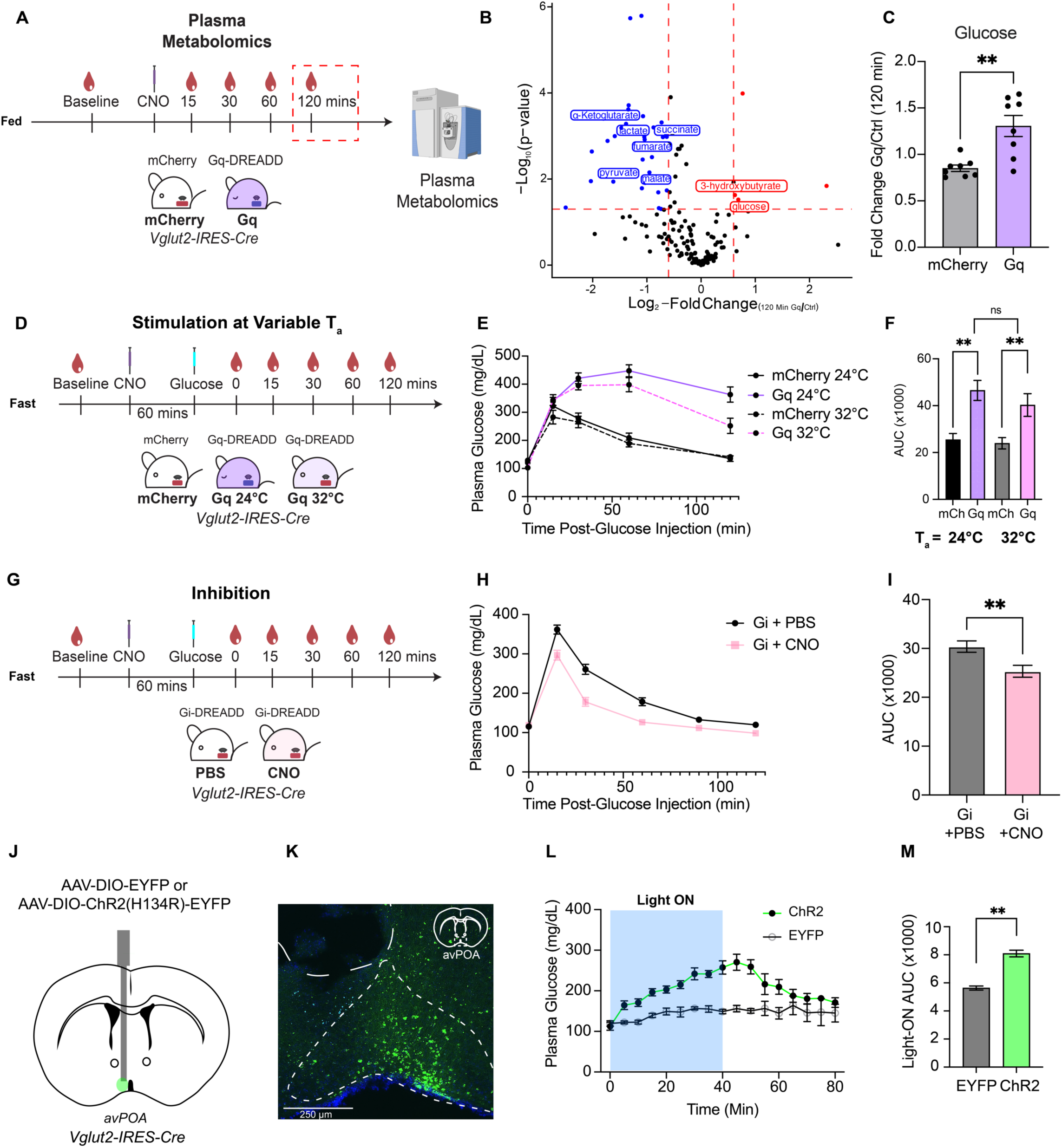
avPOA^Vglut2^ neuronal regulation of whole-body glucose homeostasis is rapid and reversible. **(A)** Schematic of blood sampling protocol for plasma metabolomics from mCherry- and Gq-DREADD-transduced *Vglut2-IRES-Cre* mice pre- and post-CNO administration. n = 8 mice (4 female, 4 male) per cohort. Blood samples were taken 0, 15, 30, 60, and 120 min post-CNO injection (1 mg/kg) for plasma separation, LC/MS, and polar metabolomics analysis. **(B)** (*Left*) Volcano plot of differentially altered metabolites 120 min post-CNO administration. Selected overrepresented and underrepresented metabolites labeled in red and blue, respectively. p-values calculated using two-tailed t-test, red lines corresponding to FDR-corrected p < 0.05, and |Log_2_-Fold change| > 0.6. **(C)** Fold-change in glucose abundance in Gq-DREADD-expressing vs. mCherry control mice at 120 min post-CNO injection. (n = 8 animals, mean ±SEM, ** = p<0.005, two-tailed t-test). **(D)** Schematic for intraperitoneal glucose tolerance test (GTT) in Gq-DREADD and mCherry control mice. Animals were fasted for 3h prior to baseline glucose measurement and CNO administration. Subsequent to a 60-minute interval following CNO administration, a baseline blood sample was taken, animals were injected with glucose (2 g/kg), and subsequent blood samples for changes in glucose levels were obtained at 15, 30, 60, and 120 min post glucose administration. **(E)** Glucose tolerance testing of mCherry (Ctrl) or Gq-DREADD-expressing animals at T_a_ = 24°C or 32°C (24°C: n = 13 mCherry, n = 14 Gq; 32°C: n = 14 mCherry, n=16 Gq *Vglut2-IRES-Cre* animals, mean ±SEM). **(F)** Area under the curve (AUC) quantification of (E) in all cohorts (** = p < 0.005, student’s two-tailed t-test, mean ±SEM). **(G)** Schematic for GTT in Gi-DREADD-transduced *Vglut2-IRES-Cre* animals. Gi-DREADD-expressing animals were injected with either PBS or CNO (10 mg/kg) prior to GTT, and blood was collected as in other GTT experiments. **(H)** Glucose tolerance testing of PBS- and CNO-injected Gi-DREADD animals (n = 8 PBS, n = 8 CNO, mean ±SEM). **(I)** AUC quantification of GTT in (H) for PBS- and CNO-injected experiments (** = p < 0.005, student’s paired two-tailed t-test, mean ±SEM). **(J)** Schematic of unilateral stereotactic administration of AAV-DIO-ChR2(H134R)-EYFP or AAV-DIO-EYFP to the avPOA of *Vglut2-IRES-Cre* mice (n=4 EYFP, n= 5 ChR2). **(K)** Representative image of (J), showing ChR2-EYFP signal in an experimental animal (Scale bar = 250 *μ*m). **(L)** Plasma glucose sampling of ChR2- and EYFP-expressing mice for 80 min at 5 min intervals (Light ON 0-40 minutes, indicated by blue shading; Light OFF 41-80 minutes, indicated by no shading, mean ±SEM). **(M)** AUC quantification of (L) (** **=** p < 0.005, Students two-tailed t-test, n = 4 EYFP, n = 5 ChR2 *Vglut2-IRES-Cre* animals).

Blood glucose levels are dynamically regulated by balancing glucose production through hepatic gluconeogenesis and glycogenolysis with tissue glucose uptake via insulin-independent and -responsive glucose transporters^17,18^. To test whether avPOA^Vglut2^ neuron stimulation resulted in reduced glucose uptake, we subjected control (mCherry-injected) and experimental (Gq-DREADD-injected) animals to a modified glucose tolerance test (GTT) following CNO stimulation. Briefly, animals were fasted for 3 hours, administered CNO, and injected intraperitoneally with a bolus of glucose (2 g/kg) 1 hour post-CNO stimulation (**Figure 2c**). Strikingly, avPOA^Vglut2^ neuron-stimulated animals were significantly hyperglycemic at the end of the test relative to their control counterparts (**Figures 2d-f**), indicating that avPOA^Vglut2^ neuron stimulation induces pronounced systemic glucose intolerance. Consistent with our previously observed effects on RER (**Figure 1f**), this glucose intolerance effect was not dependent on decreased body temperature, as raising the ambient temperature (T_a_ = 32°C) failed to restore glucose tolerance in these torpid mice (**Figure 2e-f**).

This finding led us to hypothesize that the effect of these neurons could be bidirectional, and that inhibition of these neurons may improve glucose tolerance. To test this idea, we transduced *Vglut2-IRES-Cre* animals with a Cre-dependent inhibitory Gi-DREADD (AAV-hSyn-DIO-hM4D(Gi)-mCherry). To confirm effective targeting, we fasted Gi-DREADD-transduced animals and observed that, as expected, upon CNO administration, Gi-DREADD-mediated silencing of avPOA^Vglut2^ neurons was sufficient to prevent torpor entry (**Supplementary Figures 2e, f**). To investigate the effects of avPOA^Vglut2^ neuronal silencing on glucose tolerance, we first performed a baseline GTT in which mice were pretreated with PBS and after one week of recovery, the same animals were pretreated with CNO and administered a second GTT. Remarkably, acute inhibition of avPOA^Vglut2^ neurons resulted in increased glucose tolerance, as peak glucose levels were significantly lower following CNO injection as compared to PBS (PBS Glucose_Max_ = 361.7 ± 11.4 mg/dL, CNO Glucose_Max_ = 296.9 ± 12.4 mg/dL), and remained consistently lower throughout the duration of the assay (**Figures 2h, i**). These findings suggest that avPOA^Vglut2^ neuronal activity can bidirectionally control systemic glucose tolerance.

To examine the kinetics of this induced glucose intolerance, we transduced the avPOA of *Vglut2-IRES-Cre* mice with AAV expressing either a Cre-dependent Channelrhodhopsin-2 (ChR2) (AAV-DIO-ChR2(H134R)-EYFP) or an EYFP (AAV-DIO-EYFP) control, and subsequently implanted an optic fiber above the avPOA (**Figure 2j**). After surgical recovery, we monitored plasma glucose and T_b_ over the course of 40 minutes of optogenetic blue light stimulation (16 Hz, 8.5 mW, 1s on 1s off) as well as for an additional 40 minutes post-stimulation. Optogenetic stimulation of avPOA^Vglut2^ neurons resulted in the anticipated decrease in body temperature, as well as a rapid, monotonic increase in blood glucose not observed in the EYFP controls (**Figures 2k-l, Supplementary Figure 2f**). Notably, upon light cessation, blood glucose levels and T_b_ rapidly returned to normal, demonstrating that glucose intolerance is a rapid, transient, and reversible effect of avPOA^Vglut2^ neuronal stimulation. Together, our findings identify avPOA^Vglut2^ neurons as a significant glucoregulatory population that contributes to rapid, reversible modulation of glucose homeostasis, drives a torpor-associated metabolic shift from glucose metabolism towards fatty acid catabolism upon stimulation, and bidirectionally regulates glucose tolerance upon activation or inhibition.

### avPOA^Vglut2^ neuronal activation drives insulin resistance and reduces glucose uptake

Dysregulation of glucose homeostasis is often secondary to either reduced insulin secretion or reduced insulin action on peripheral tissues in response to glucose challenge^17^. To test whether the observed avPOA^Vglut2^ neuron-induced dysregulation of glucose clearance was a consequence of reduced insulin secretion, we assessed circulating insulin levels in the context of our GTT experimental paradigm. Remarkably, we observed no significant differences in insulin levels in CNO-stimulated Gq-DREADD-expressing animals relative to mCherry controls (**Supplementary Figure 3a**), suggesting that glucose-stimulated insulin secretion (GSIS) is unaffected by avPOA^Vglut2^ neuronal stimulation. These findings were further confirmed by monitoring GSIS in both Gq-DREADD-expressing and control mice in the context of a hyperglycemic clamp^19^ (**Supplementary Figure 3b**). Indeed, even the elevated insulin levels observed in Gq-DREADD-expressing mice could not normalize plasma glucose in these animals following CNO-stimulation, suggesting that avPOA^Vglut2^ neuron stimulation induces systemic insulin insensitivity.

To further investigate this avPOA^Vglut2^ neuronal stimulation-induced insulin insensitivity, we performed hyperinsulinemic-euglycemic clamp studies in Gq-DREADD-expressing *Vglut2-IRES-Cre* mice and corresponding mCherry control animals following CNO stimulation (**Figure 3a**). Briefly, in both Gq-DREADD- and mCherry-expressing animals, we induced hyperinsulinemia via constant insulin infusion while concomitant glucose infusion at a variable rate was used to maintain euglycemia^20^. Importantly, all animals were partially restrained during these studies for the purposes of blood sampling. This afforded us an ability to control for the reduced locomotor activity of avPOA^Vglut2^ neuron-stimulated mice as compared to their unstimulated control counterparts (see **Methods**).

**Figure 3:**
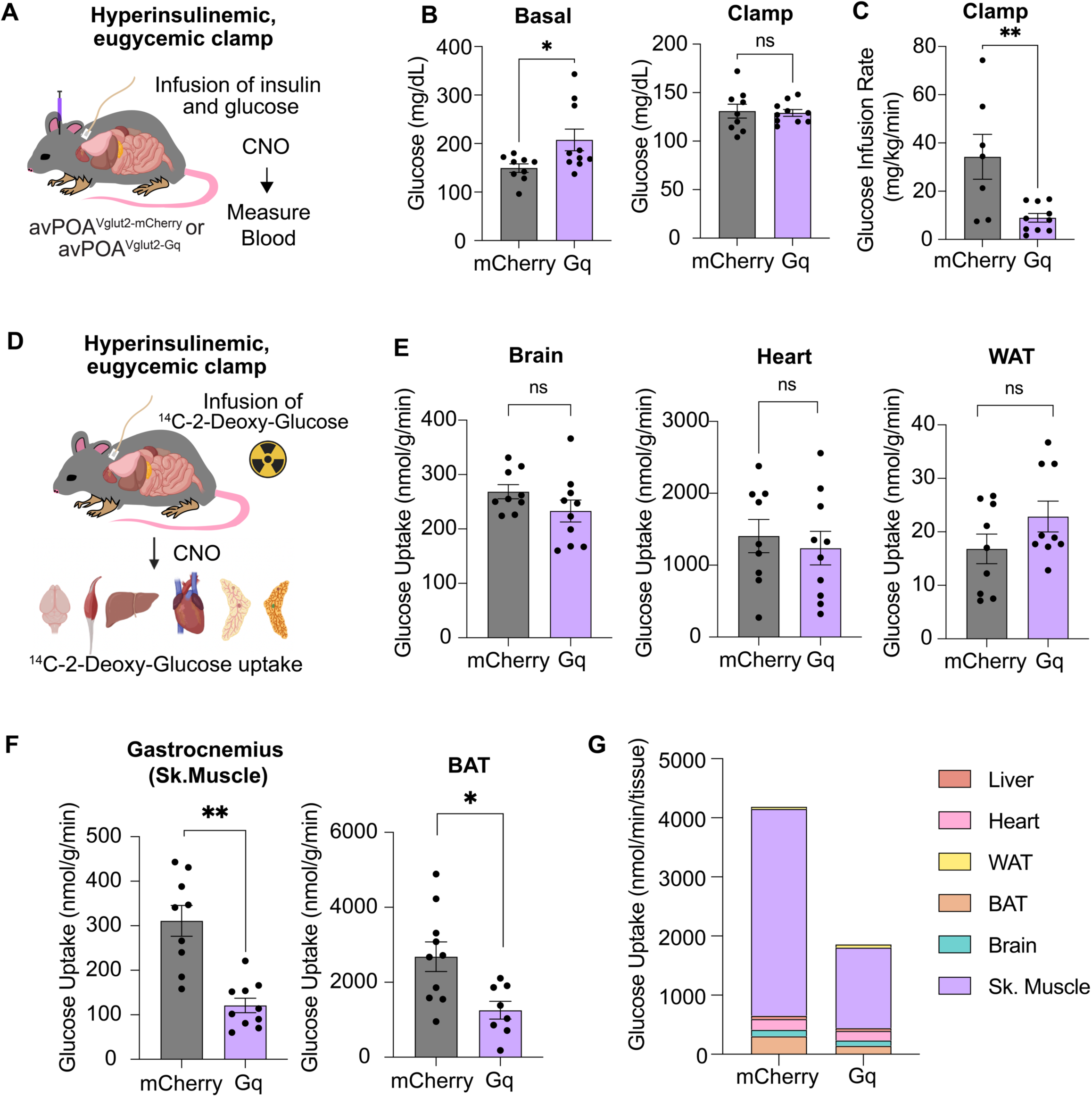
avPOA ^Vglut2^ neurons coordinate whole-body glucose hypometabolism and organ-specific defects in glucose uptake. **(A)** Schematic for hyperinsulinemic-euglycemic clamp and tissue-specific glucose uptake measurements. Gq-DREADD- or mCherry-transduced animals were implanted with indwelling catheters, mildly restrained, and subjected to continuous infusion of glucose, insulin, ^3^H-glucose, and ^14^C-2-deoxyglucose over the course of the two-hour experiment. Glucose levels were maintained at approximately 120 mg/dL to ensure clamp effectiveness. Animals were sacrificed, and the displayed tissues were taken for radioactive ^14^C-2-Deoxyglucose-6-Phosphate uptake analysis. **(B)** Glucose levels 60 min after CNO stimulation (Basal) were significantly elevated in Gq-DREADD-expressing animals as compared to control animals (*\* =* p<0.05, student’s two-tailed t-test, mCherry *=* 149.4 ± 8.8 mg/dL; Gq *=* 207.5 ± 22.2 mg/dL, mean ±SEM, n= 9 mCherry, n=10 Gq *Vglut2-IRES-Cre* animals). Conversely, during the hyperinsulinemic clamp, Gq-DREADD- and mCherry-expressing animals maintained nearly identical levels plasma glucose (Gq: 130.9 ± 7.2 mg/dL, mCherry: 129.1 ± 3.4 mg/dL), indicating successful clamping of plasma glucose. **(C)** Glucose Infusion Rate (GIR) of control (mCherry) and Gq-DREADD-transduced animals throughout the 120-minute duration of the experiment (** = p < 0.005, student’s two-tailed t-test, mean ±SEM). **(D)** Organs in which avPOA stimulation had no significant effect on insulin-stimulated glucose uptake (ns = p > 0.05, student’s two-tailed t-test, mean ±SEM). **(E)** Gastrocnemius and BAT exhibited reductions of 61.1% and 55.3% in ^14^C-2-DG-6-Phosphate uptake (Glucose Uptake), respectively, between Gq-DREADD- and mCherry-expressing animals (* = p < 0.05, ** = p< 0.005, student’s two-tailed t-test, mean ±SEM). **(F)** Calculated mass-dependent glucose uptake in each of the six examined organs based on average tissue weights of control and Gq-DREADD-expressing animals and tissue-specific glucose uptake rates (Mass-Dependent Uptake (nmol/min) = Average Tissue Mass (g) * Glucose Uptake Rate (nmol/min/g tissue)).

Although euglycemia was achieved in both the Gq-DREADD-expressing and control cohorts, the glucose-infusion rate (GIR) to maintain euglycemia during the insulin clamp experiments in Gq-DREADD-expressing animals was significantly lower than that of controls, indicating reduced insulin sensitivity (**Figures 3b, c)**. Moreover, while hepatic glucose production during the insulin clamp was unaffected by avPOA^Vglut2^ neuronal stimulation, both whole-body glucose turnover and glycolysis were dramatically reduced (**Supplementary Figures 3f-i**), consistent with our observed metabolomics results in which circulating glycolytic and TCA cycle metabolites were reduced by avPOA^Vglut2^ neuron stimulation. Taken together, our findings show that avPOA^Vglut2^ neuronal activation induces systemic insulin resistance in mice, thereby contributing to reduced glucose metabolism in these animals.

### avPOA^Vglut2^ neurons control organ-specific changes in glucose uptake

To characterize the individual organs responsible for systemic insulin resistance, a bolus of the non-metabolizable glucose analogue, ^14^C-2-deoxy-glucose (^14^C-2DG), was intraperitonelly injected during the hyperinsulinemic-euglycemic clamp experiments. At the conclusion of the experiments, animals were sacrificed, and liver, brain, white adipose tissue (WAT), brown adipose tissue (BAT), heart, and gastrocnemius muscle were collected for biochemical analysis of their ^14^C-2DG-6-phosphate content. Scintillation counting revealed no significant changes in the insulin-stimulated glucose uptake rate of heart, WAT, and brain. However, there was a marked reduction in glucose uptake in both gastrocnemius muscle and BAT from avPOA^Vglut2^ neuron-stimulated animals (**Figures 3d, e**), indicating organ-specific changes in insulin action and glucose metabolism.

We calculated total glucose uptake in the presence or absence of avPOA^Vglut2^ neuronal stimulation for each of the six tested tissues based on our measured rates of glucose uptake and tissue weights. Consistent with prior findings that skeletal muscle accounts for ∼80% of post-prandial glucose utilization, and up to 75% of glucose disposal upon infusion^21^, this analysis revealed that diminished skeletal muscle glucose uptake accounts for a large fraction of the total glucose utilization deficit observed in avPOA^Vglut2^ neuron-stimulated animals (**Figure 3f**), suggesting that effects on skeletal muscle are the major contributor to the avPOA^Vglut2^ neuron-driven impairment of insulin sensitivity. Importantly, we confirmed that this decrease in skeletal muscle glucose uptake was not dependent on body temperature changes (**Figures 4a, b**). In addition, comparative phospho-proteomic analysis of gastrocnemius samples from avPOA^Vglut2^ neuron-stimulated and control animals provided evidence of avPOA^Vglut2^ neuron-driven suppression of skeletal muscle insulin signaling (**Supplementary Figure 4c-f**). Given these findings, we focused our subsequent investigation of the peripheral effects of avPOA^Vglut2^ neuron activation on skeletal muscle.

**Figure 4:**
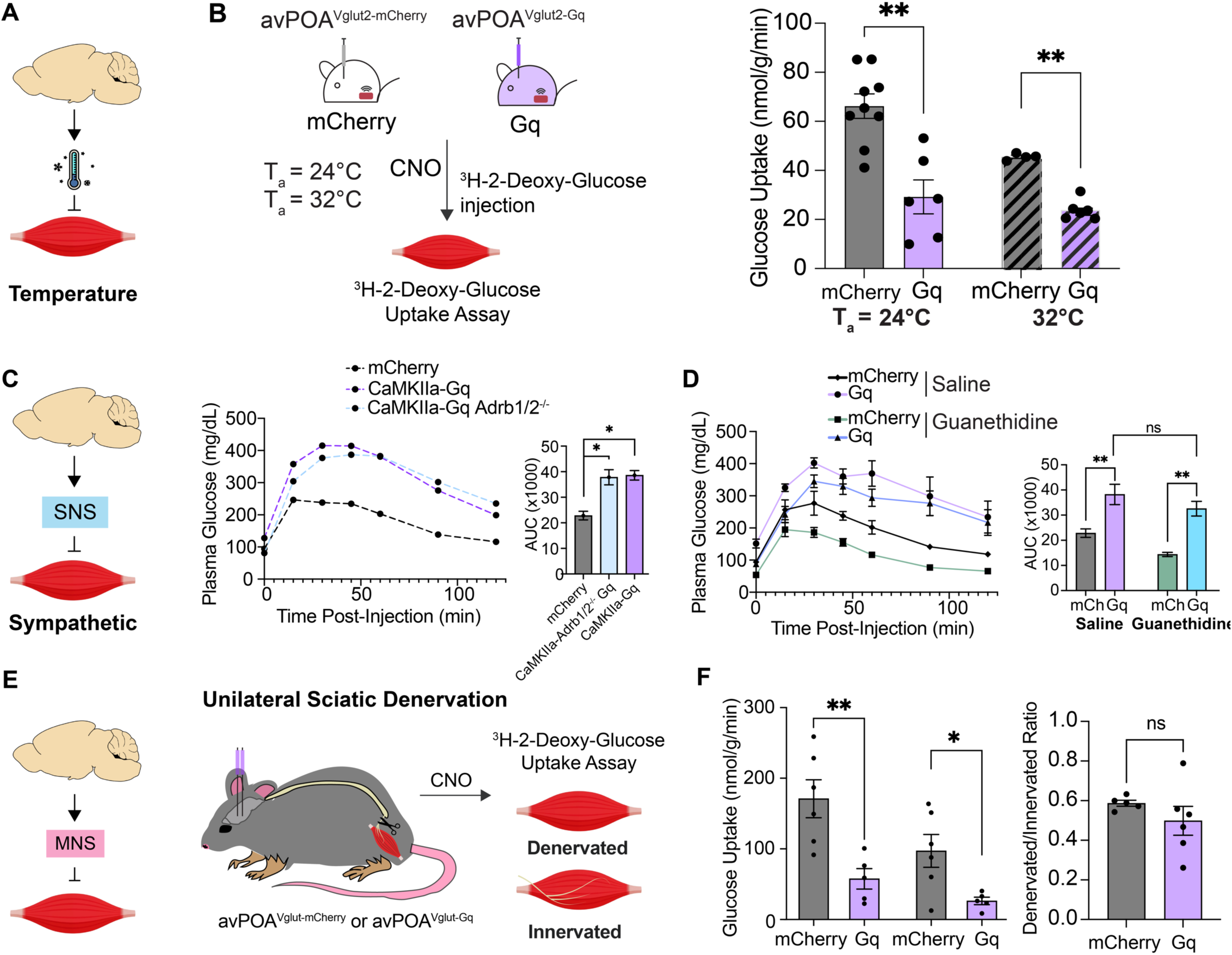
Determination of peripheral effectors of avPOA^Vglut2^-induced skeletal muscle insulin resistance. **(A)** Schematic for potential temperature-regulated effects of avPOA^Vglut2^ neuronal stimulation on skeletal muscle glucose uptake. **(B)** (*Left*) Schematic for radioactive glucose (^3^H-2-deoxyglucose-6 phosphate) uptake assay in *Vglut2-IRES-Cre* animals injected with either Cre-dependent mCherry (avPOA^Vglut2-mCherry^) or Gq-DREADD (avPOA^Vglut2-Gq^), then pretreated with CNO and subjected to radioactive glucose uptake assay at T_a_ = 24°C or 32°C. (*Right)* results of Radioactive glucose (^3^H-2-deoxyglucose-6 phosphate) uptake assay in gastrocnemius muscles of control (avPOA^Vglut2-mCherry^) or Gq-DREADD (avPOA^Vglut2-Gq^) mice housed at T_a_ =24°C or 32°C (** = p < 0.005, student’s two-tailed t-test, mean ±SEM, n = 9 mCherry 24°C, n = 6 Gq 24°C, n = 4 mCherry 32°C, n=6 Gq 32°C *Vglut2-IRES-Cre* animals). **(C)** Glucose tolerance testing of control mCherry-expressing mice (mCherry), mice transduced with AAV-CaMKIIa-hM3D(Gq)-mCherry (CaMKIIa-Gq), and CaMKIIa-Gq-transduced *Adrb1/2^-/-^*mice (CaMKIIa-Gq Adrb1/2^-/-^). AUC calculation shown on right (* = p < 0.05, student’s two-tailed t-test, mean ±SEM, n = 6 Vglut-mCherry, n = 6 CaMKIIa-Gq, n = 4 CaMKIIa-Gq Adrb1/2^-/-^ animals). **(D)** Pair-matched glucose tolerance tests in animals injected with saline, CNO (1 mg/kg), Guanethidine (50 mg/kg), or Guanethidine and CNO (ns = p > 0.05, ** = p < 0.005, paired student’s two-tailed t-test, mean ±SEM, n = 6 mice per condition). **(E)** Schematic of sciatic nerve denervation paradigm to test the requirement for motor innervation for avPOA-induced skeletal muscle insulin resistance. Unilateral denervation allows for a matched innervated internal control. **(F)** Radioactive glucose uptake assay in innervated or denervated gastrocnemius muscle in CNO-treated control mCherry- and Gq-DREADD-expressing animals (mCherry innervated = 171.0±26.9, Gq innervated = 57.6±14.5, mCherry denervated = 97.1±23.2, Gq denervated = 26.3±5.3 nmol/g/min). (*Right*) Ratio of denervated:innervated gastrocnemius glucose uptake in mCherry animals (59.9±1.5%) and Gq animals (49.0±7.3%).

### avPOA^Vglut2^ neuronal control of muscle glucose uptake occurs independent of muscle innervation

We next sought to determine the manner by which avPOA^Vglut2^ neurons convey glucoregulatory signals to the periphery (**Figure 4a**). In this regard, skeletal muscle is innervated by both sympathetic nervous system (SNS) and motor nervous system (MNS) fibers^22^, and sympathetic activation has been previously shown to enhance skeletal muscle glucose uptake via the β-adrenergic signaling, primarily through β2-adrenergic receptors^23^. We therefore employed a genetic approach to test the requirement for β-adrenergic sympathetic signaling in avPOA^Vglut2^ neuron-driven changes in glucose tolerance, utilizing mice lacking β_1_- and β_2_-adrenergic receptors (β_1/2_-KO). For these studies, a *CaMKIIa* promoter-driven AAV (AAV-CaMKIIa-hM3D(Gq)-mCherry) was used to achieve selective Gq-DREADD expression in excitatory avPOA neurons of β_1/2_-KO and wild-type control mice. To ensure that our results with this *CaMKIIa* approach were consistent with our *Vglut2-IRES-Cre* experiments, we compared expression patterns and GTT results of *CaMKIIa*-Gq-injected animals with those of Cre-dependent *Vglut2-IRES-Cre* animals and found no significant differences in expression nor glucose intolerance between cohorts (**Supplementary Figure 4g**). Following CNO stimulation, we performed a GTT assay on these animals, which revealed no significant differences in CNO-stimulated glucose intolerance between Gq-DREADD-expressing β_1/2_-KO and wild-type animals (**Figure 4c**), suggesting that β_1/2_ adrenergic receptors were not required for avPOA^Vglut2^ glucoregulatory signaling. As an orthogonal approach, we also performed chemical sympathetic inhibition in Gq-DREADD-expressing *Vglut2-IRES-Cre* mice via the sympatholytic drug guanethidine, which enters noradrenergic terminals and displaces norepinephrine from synaptic vesicles to suppress norepinephrine release^24^. As expected, given its vasodilatory properties, guanethidine treatment alone increased baseline glucose tolerance compared with PBS. However, despite SNS inhibition by guanethidine, avPOA^Vglut2^ activation still induced glucose intolerance (**Figure 4d**). Taken together, these genetic and pharmacological studies strongly suggest that SNS activity is dispensable for avPOA^Vglut2^ neuron-driven glucose intolerance.

Motor innervation and movement also serve as well-appreciated regulators of glucose metabolism^18^. To test the role of motor innervation on glucose uptake during avPOA^Vglut2^ neuronal stimulation, we performed unilateral sciatic nerve denervations in mCherry control and Gq-DREADD-transduced *Vglut2-IRES-Cre* mice^25^. This approach allowed us to compare, within the same animal, the effects of avPOA^Vglut2^ neuron stimulation on muscle glucose uptake in the innervated and the denervated limb (**Figure 4e**). Unilaterally denervated animals were injected with CNO 1 hour prior to administration of the nonmetabolizable radioactive glucose analog ^3^H-2-Deoxyglucose. The animals were then monitored for changes in blood glucose and radioactive substrate availability for 30 minutes, sacrificed, and their ipsi- and contra-lateral gastrocnemius muscles processed for scintillation counting of ^3^H-2-DG-6-Phosphate. As expected, denervated gastrocnemius muscle exhibited reduced glucose uptake relative to the contralateral innervated muscle under basal conditions (**Figure 4f**). Importantly, both innervated and denervated muscles exhibited similarly pronounced reductions in glucose uptake following CNO-mediated avPOA^Vglut2^ neuronal stimulation, with no significant differences in the ratio of CNO-stimulated to baseline glucose uptake in the innervated and denervated muscle (**Figure 4f**). Taken together, these results demonstrate that sympathetic innervation and direct motor innervation are dispensable for avPOA^Vglut2^ neuron-driven changes in muscle glucose uptake.

### Corticosterone contributes to the avPOA^Vglut2^-induced reduction of muscle glucose uptake

Given the apparent dispensability of direct muscle innervation for avPOA^Vglut2^ neuron-to-skeletal muscle signaling, we examined the potential involvement of known circulating glucoregulatory hormones, assessing insulin, glucagon, leptin, and corticosterone plasma levels in Gq-DREADD- and mCherry-transduced animals by ELISA following treatment with CNO (**Figure 5a**). While no significant differences in insulin, glucagon, or leptin levels were associated with avPOA^Vglut2^ neuronal stimulation (**Figure 5a**), we observed a ∼2.5-fold increase in plasma corticosterone levels in Gq-DREADD-expressing animals up to 2 hours post-CNO stimulation, as compared with baseline (**Figure 5a**). The increase in corticosterone tracked well with the observed increase in glucose levels (**Figure 5b**), suggesting a potential role for circulating corticosterone levels in avPOA^Vglut2^ neuron-induced glucose uptake changes.

**Figure 5:**
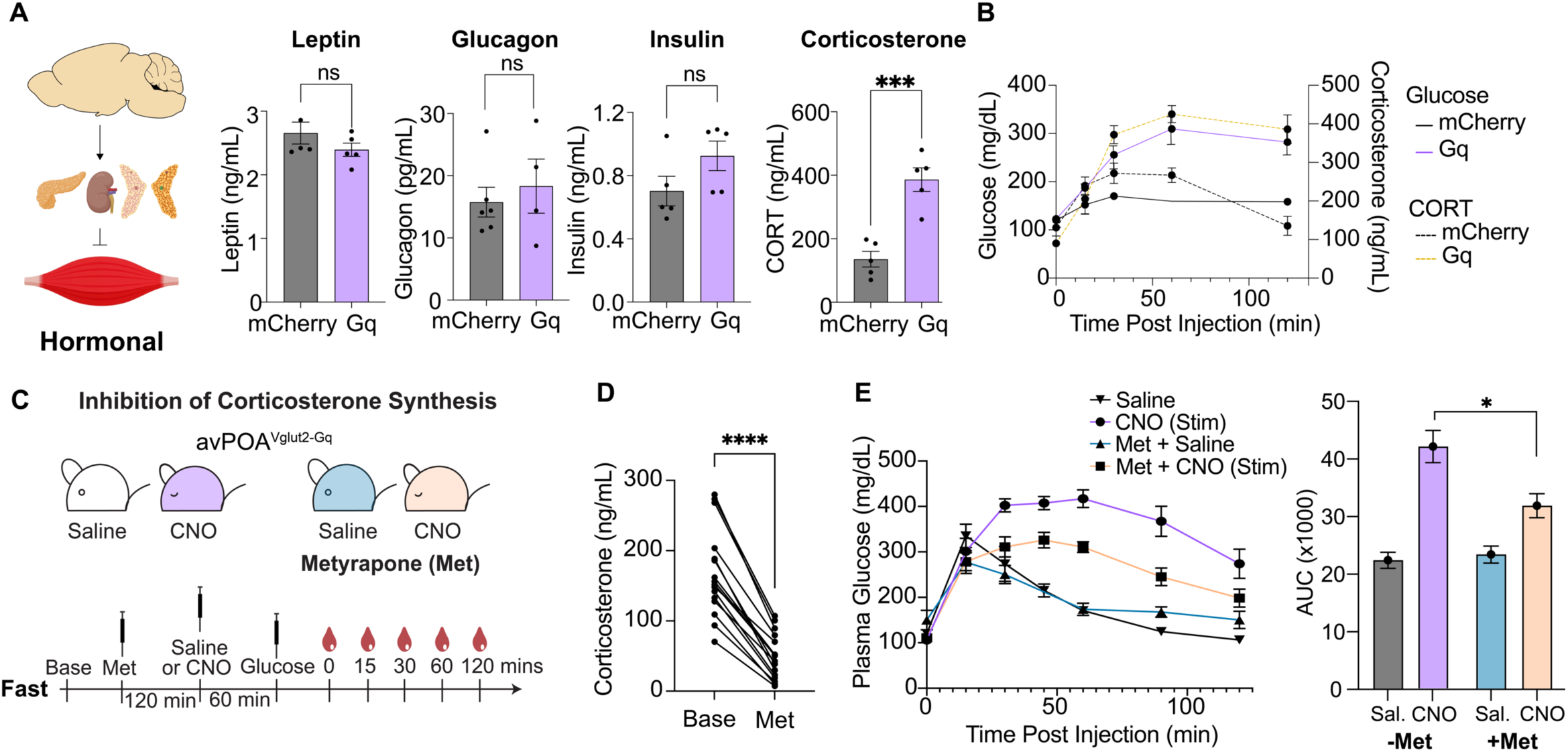
avPOA-induced insulin intolerance requires corticosterone. **(A)** Concentrations of plasma leptin, glucagon, insulin, and corticosterone 60 minutes following CNO administration in fed mCherry- and Gq-DREADD-expressing animals. (Leptin: n = 5 mCherry, n = 5 Gq; Glucagon: n= 5 mCherry, n = 6 Gq; Insulin: n = 5 mCherry, n = 5 Gq; Corticosterone: n = 5 mCherry, n = 7 Gq animals, ** = p < 0.005). **(B)** Longitudinal monitoring of glucose and corticosterone in fed mCherry- and Gq-expressing mice injected with 1 mg/kg CNO. Blood plasma taken 0, 15, 30, 60, and 120 minutes post-injection (n = 7 Gq, n = 5 mCherry Vglut2-IRES-Cre animals). **(C)** Schematic for metyrapone treatment during glucose tolerance testing. Animals were fasted for 120 minutes before baseline blood glucose levels were taken. Animals were then injected with 50 mg/kg metyrapone, and CNO was administered 2 hours following metyrapone injection. Subsequently, the GTT proceeded as previously described. **(D)** Corticosterone ELISA on blood sampled from animals prior to metyrapone injection (Base) or 2 hours post-injection (Met) (**** = p<0.00005, student’s two-tailed t-test, mean ±SEM, n = 6 samples per condition). (**E)** Glucose tolerance test of Gq-DREADD-transduced *Vglut2-IRES-Cre* animals injected with saline, CNO (1 mg/kg), saline + metyrapone (50 mg/kg), or CNO (1 mg/kg) + metyrapone (50 mg/kg). (*Right*) AUC quantitation of results (* = p < 0.05, student’s two=tailed t-test, mean ±SEM, n = 6 *Vglut2-IRES-Cre* animals).

To directly test the contribution of corticosterone to avPOA^Vglut2^ neuron-induced glucose intolerance, we utilized metyrapone, an inhibitor of corticosterone synthesis (**Figure 5c**)^26^. Baseline blood samples from unstimulated Gq-DREADD-expressing *Vglut2-IRES-Cre* mice confirmed that metyrapone administration significantly decreased serum corticosterone levels relative to saline-treated controls (**Figure 5d**). Subsequent GTT assays showed that metyrapone treatment significantly attenuated CNO-induced glucose intolerance in Gq-DREADD-expressing animals (**Figures 5e**). Although incomplete, this reversal of glucose intolerance in metyrapone-treated animals identifies adrenal-derived corticosterone as a major regulator of glucose homeostasis downstream of avPOA^Vglut2^ neuronal activation.

### avPOA^PACAP^ neuronal activity is sufficient to drive avPOA glucoregulatory effects

Having outlined this POA-to-muscle glucoregulatory signaling axis, we sought to better define the neuronal population within the POA that underlies these glucoregulatory effects. To begin to address this question, we performed axon terminal and monosynaptic anterograde mapping of downstream avPOA^Vglut2^ target regions (**Figure 6a, Supplementary Figures 5a, b**)^27^. Both mapping approaches revealed the dorsomedial hypothalamus (DMH) and Raphe Pallidus (RPa) as densely innervated by avPOA^Vglut2^ neurons. These regions have been heavily implicated in control of metabolic rate and thermogenesis^28,29^. To test their role in avPOA^Vglut2^ neuron-driven regulation of glucose homeostasis, we optogenetically stimulated axonal terminals of avPOA^Vglut2^ neurons transduced with Cre-dependent ChR2 in either the DMH or RPa and analyzed the effects on blood glucose tolerance. Despite dense axonal labeling in the RPa, stimulation of avPOA^Vglut2^ neuron terminals in this region failed to induce glucose intolerance (**Figure 6c**). By contrast, stimulation of DMH-projecting neurons fully phenocopied the glucose intolerance induced by avPOA soma stimulation (**Figure 6c**), indicating that excitatory avPOA->DMH projections are sufficient to drive acute glucose intolerance.

**Figure 6:**
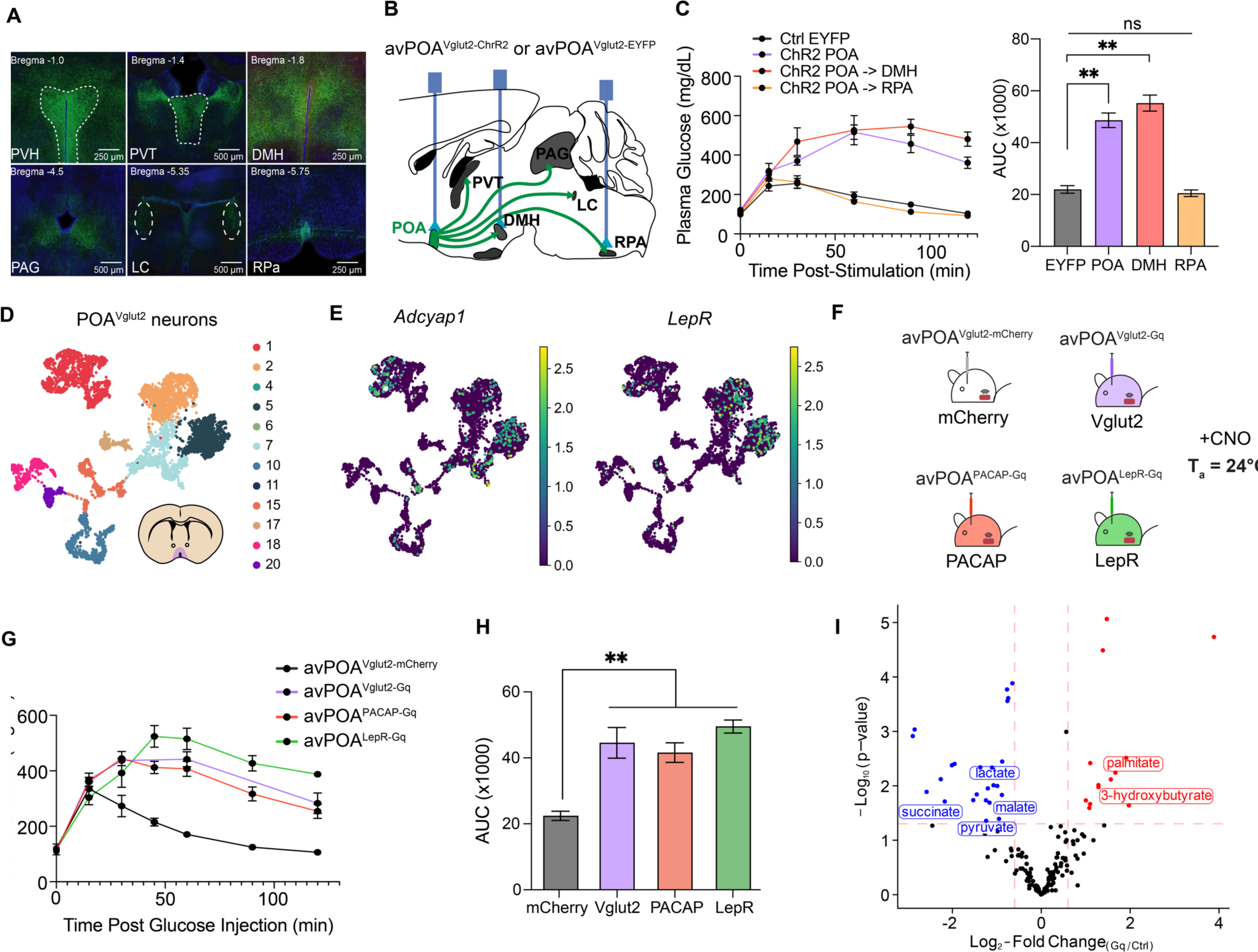
avPOA^PACAP^ and avPOA^LepR^ neuronal subsets function as regulators of metabolic flexibility. **(A)** Immunostaining of axonal fibers in *Vglut2-IRES-Cre* mice transduced with Gq-DREADD (AAV-hSyn-DIO-hM3D(Gq)-mCherry). Axonal projections were identified in a variety of downstream regions, including the paraventricular thalamus (PVT), paraventricular hypothalamus (PVH), dorsomedial hypothalamus (DMH), periacqueductal grey (PAG), locus coeruleus (LC), and raphe pallidus (RPa). (**B)** Optogenetic branch stimulation of hindbrain-projecting avPOA^Vglut2^ neurons. Blue bars correspond to regions targeted for optogenetic stimulation via cannula implantation and subsequent exposure to 455 nm blue light. **(C)** (*Left)* Glucose tolerance testing of *Vglut2-IRES-Cre* animals transduced with a Cre-dependent ChR2 (AAV-DIO-ChR2(H134R)-EYFP) or EYFP (AAV-DIO-EYFP) with optic fibers placed unilaterally above either the POA, POA→DMH-projecting neurons, or POA→RPA-projecting neurons. *(Right)* AUC of GTT for each cohort described (ns = p > 0.05, **\*\* =** p < 0.005, student’s two-tailed t-test, mean ±SEM, n = 4 EYFP, n = 4 POA, n = 3 POA→DMH, n = 3 POA→RPa *Vglut2-IRES-Cre* animals). **(D)** UMAP plot and Leiden clustering of all avPOA^Vglut2^ neurons identified in Hrvatin et al., 2020. **(E)** UMAP plot of avPOA^Vglut2^ neurons colored by *Adcyap1* and *LepR* expression. **(F)** Schematic showing four groups of animals: mCherry (mCherry-transduced *Vglut2-IRES-Cre,* n = 11), Vglut2 (Gq-DREADD transduced *Vglut2-IRES-Cre*) (n=10), PACAP (Gq-DREADD-transduced *PACAP-2A-Cre*) (n=11), and LepR (Gq-DREADD-transduced *LepR-IRES-Cre* (n=3) animals injected with 1 mg/kg CNO. **(G)** Glucose tolerance test of mCherry (avPOA^Vglut2-mCherry^), Vglut2 (avPOA^Vglut2-Gq^), PACAP (avPOA^PACAP-Gq^), and LepR (avPOA^LepR-Gq^), animals from (F). **(H)** AUC analysis for (G) (** = p < 0.05, student’s two-tailed t-test, mean ±SEM, n=7 avPOA^Vglut2-mCherry^, n = 10 avPOA^Vglut2-Gq^, n = 11 avPOA^PACAP-Gq^, n = 3 avPOA^LepR-Gq^ animals). (**I)** Volcano Plot of 120-minute polar metabolomics of *PACAP-2A-Cre* animals transduced with a Cre-dependent Gq-DREADD and injected with either PBS or CNO (1 mg/kg). Animals were treated as in Figures 2a-c. Selected overrepresented and underrepresented metabolites labeled in red and blue, respectively. p-values calculated using two-tailed t-test; red lines corresponding to p-value and Log_2_-Fold changes set at p < 0.05 and |Log_2_-Fold change| > 0.6.

In addition to the anatomical heterogeneity of its projections, the avPOA glutamatergic neuronal population is comprised of molecularly diverse neuronal subpopulations that could mediate distinct effects on physiology^30^. In this regard, we initially focused on Adcyap1 (PACAP)- and Leptin Receptor (LepR)-expressing neurons given the established involvement of these two largely excitatory POA neuronal subpopulations in metabolic and thermoregulatory programs^13,31^. Integrated analysis of our prior single-nucleus transcriptomic data^11^, showed that *Adcyap1*+ neurons largely constitute a subset of POA glutamatergic neurons present throughout the anatomical length of the POA, whereas LepR+ neurons form a distinct but molecularly overlapping population with *Adcyap1* neurons located primarily in the anterior portion of the POA (**Figures 6d, e**). To determine whether chemogenetic stimulation of one or both of these subpopulations was sufficient to give rise to avPOA^Vglut2^-associated glucoregulatory effects, we stereotactically introduced our Cre-dependent AAV expressing Gq-DREADD into the avPOA of *Adcyap1-2A-Cre* (avPOA^PACAP^) or *Lepr-IRES-Cre* (avPOA^LepR^) animals. Subsequent chemogenetic stimulation of both avPOA^PACAP^ and avPOA^LepR^ neurons fully phenocopied the glucose intolerance observed upon avPOA^Vglut2^ neuronal stimulation (**Figures 6f-h**), demonstrating that either avPOA^PACAP^ or avPOA^LepR^ neuronal stimulation is sufficient to give rise to the glucoregulatory effects of the broader avPOA^Vglut2^ population.

Given these results implicating PACAP^+^ avPOA neurons in the regulation of blood glucose homeostasis, we sought to demonstrate their importance in regulating organismal metabolic flexibility. To this end, we repeated our prior polar metabolomics approach done in *Vglut2-IRES-Cre* animals with *PACAP-2A-Cre* mice that were stereotactically transduced with Cre-dependent mCherry or Gq-DREADD AAVs, and upon recovery subjecting these animals to blood collection and metabolic profiling following CNO stimulation. Consistent with the results observed under avPOA^Vglut2^ neuronal stimulation, we observed significant decreases in the abundance of major glycolytic and citric acid cycle metabolites when comparing avPOA^PACAP^ Gq-DREADD-stimulated and control avPOA^mCherry^ animals 120 minutes after CNO administration, including pyruvate, citrate, succinate, lactate, and malate (**Figure 6i**). Moreover, consistent with increased reliance on fatty acid oxidation for fuel, we observed significant increases in β-hydroxybutyrate in Gq-DREADD-stimulated avPOA^PACAP^ animals. There was also a modest but non-significant increase in glucose abundance in these same animals 120 minutes post-stimulation. Analysis of earlier time points revealed a significant increase in blood glucose in Gq-DREADD- vs. mCherry-expressing avPOA^PACAP^ animals at the 60-minute time point (**Supplementary Figures 5d, e**), consistent both with our results in *Vglut2-IRES-Cre* animals and a model in which avPOA^PACAP^ neurons directly regulate glucose tolerance. Finally, we observed significant increases in the abundance of palmitate and α-lipoic acid at this time point, indicating an increased mobilization of fat reserves to provide fuel via FAO during avPOA^PACAP^ neuron stimulation. These results, coupled with our previous metabolomics experiments in *Vglut2-IRES-Cre* animals, confirm that avPOA^Vglut2^ neurons are powerful regulators of metabolic flexibility, and that avPOA^PACAP^ neurons are a Vglut2^+^ subtype sufficient to drive this metabolic switch from carbohydrate to fatty acid catabolism.

## Discussion

The ability to dynamically switch between metabolic fuel sources in the face of changing nutrient availability and energetic demands is an essential feature of metabolic regulation. Classically, this metabolic flexibility was understood to be driven mainly by circulating hormonal factors, such as insulin and glucagon, with less known regarding the role of the central nervous system in the control of whole-body and organ-specific fuel usage. Here, we focus on a population of neurons associated with torpor and heat-defense to probe the mechanisms by which CNS signals drive rapid systemic metabolic reprogramming in response to environmental challenge. Surprisingly, we find that avPOA^Vglut2^ neuronal activation not only regulates whole-body metabolic flexibility, but also controls fuel uptake and catabolism in an organ-specific manner. Through a variety of interventions, we show that these effects are mediated via the induction of acute insulin insensitivity in skeletal muscle and brown adipose tissue. In skeletal muscle, this effect does not require sympathetic nervous system or motor neuron input. Rather, the glucoregulatory effects of avPOA^Vglut2^ neurons appear to be mediated, at least in part by corticosterone. Strikingly, acute silencing of this avPOA^Vglut2^ population was also found to improve glucose tolerance, demonstrating that these neurons exert bidirectional control over systemic glucose homeostasis and metabolic flexibility. While circuits regulating glucose homeostasis and the counter-regulatory response to fasting have been described in other brain regions^6,7,32^, it is notable that the effects of avPOA^Vglut2^ activation and silencing on plasma glucose levels exceed those of other known glucoregulatory circuits, suggesting that avPOA^Vglut2^ neurons may be among the most potent central regulators of glucose homeostasis identified to date.

Torpor is the primary physiological paradigm in which avPOA^Vglut2^ neuron function has been described, and represents a dramatic challenge to normal thermal and metabolic homeostasis^11,12^. In addition to their glucoregulatory capacities, activation of these neurons results in profound hypothermia and hypometabolism, which are understood to be protective adaptations for organismal survival^2^. One core aspect of this hypometabolism has been hypothesized to be the preservation of glucose for the brain via reduction in peripheral organ glucose uptake. However, until recently, it was unclear whether these neurons played an active role in shaping specific, regulated aspects of this hypometabolic state, or if torpid metabolism was secondary to decreases in T_b_. Notably, a recent study examining a molecularly distinct subpopulation of avPOA^Vglut2^ neurons expressing the neuropeptide QRFP (avPOA^QRFP^ neurons) found that activation of this subpopulation results in both torpor-induction and impaired glucoregulation, similar to avPOA^Vglut2^ neurons described in this work^33^. However, unlike our findings, the researchers reported that avPOA^QRFP^ glucoregulation is secondary to body temperature decreases. Our work shows that this is not a general principle of avPOA^Vglut2^ neuron-mediated glucoregulation. Metabolic cage, GTT, and tissue-specific glucose uptake experiments performed at elevated T_a_, where animal T_b_ minimally decreases during avPOA^Vglut2^ neuronal stimulation, all failed to ameliorate the observed glucose intolerance and uptake defects. These divergent results suggest that there may be distinct subpopulations of avPOA^Vglut2^ neurons which have temperature-dependent and -independent effects on glucose homeostasis.

Our data suggest a model in which activation of avPOA^Vglut2/PACAP^ neurons contributes to the reduction of glucose uptake in two of the primary consumers of glucose – skeletal muscle and brown adipose tissue – thereby conserving glucose for uptake by the brain. Reduction of glucose uptake in these tissues, coupled with peripheral reliance on fat stores and ketone bodies, may allow for the stabilization of glucose levels such that the animal has sufficient available energy stores to forage for food upon exit from torpor. While our results agree with studies of overnight fasting that demonstrate relative insulin resistance and glucose intolerance compared to mice with ad-libitum access to food^34^, several glucoregulatory modalities (including changes to insulin and glucagon levels) regulate glucose levels during fasting. Thus, further studies will be needed to better understand the physiological benefit of avPOA^Vglut2/PACAP^-mediated glucoregulation during torpor. Moreover, avPOA^PACAP^ neurons have been implicated in glucoregulation during warm challenge^35^, and are involved in diverse behaviors, including sleep, thirst, and cold-defense^31–37^. Future studies should examine the extent to which these neurons regulate metabolic flexibility across these diverse physiological states.

An interesting aspect unaddressed by this study is what sensory modalities serve to engage the activity of these neurons. Do these neurons respond directly to changes in circulating hormone levels or metabolite abundance to trigger glucoregulatory metabolic shifts? Are there ascending interoceptive neurocircuits that signal whole-body or tissue-specific nutrient requirements to avPOA^Vglut2/PACAP^ neurons? Future studies will be required to determine the extent to which the glucoregulatory effects of these neurons impact mouse physiology during these and other phenomenologically diverse states, and which sensory modalities trigger their activation.

Intriguingly, this hypothalamic circuitry is largely conserved in non-torpid mammals, including humans. Indeed, activation of avPOA^Vglut2^ neurons has been shown to induce torpor-like hypometabolic state in the rat, a non-torpid animal^12^. Moreover, single-cell profiling of avPOA subpopulations in non-human primates, such as marmosets, has revealed the presence of avPOA^Vglut2/PACAP^ neurons in the preoptic area of these animals (**Supplementary Figure 6**)^38,39^. While the function of these neurons in marmoset thermo- and gluco-regulation remains unexplored, the anatomical and molecular conservation of these neurons in the preoptic area suggests that they could have similar roles in regulating marmoset metabolic flexibility and metabolic homeostasis. In humans, metabolic inflexibility is primarily understood within pathological states such as diabetes, obesity, and metabolic syndromes, in which desensitization mechanisms render peripheral tissues unable to respond to physiological levels of glucoregulatory hormones such as insulin and leptin^21,36^. However, our findings raise the possibility that such metabolic states can also be initiated or augmented by top-down hypothalamic signaling. Future investigation will determine whether systematic changes in the activity of these circuits over time predispose individuals to these disorders and facilitate the development of metabolic pathologies.

## Methods and Materials

### Mice

Animal experiments were approved by the National Institutes of Health, Massachusetts Institute of Technology’s Committee on Animal Care (CAC), and Beth Israel Deaconess Medical Center’s IACUC following ethical guidelines described in the US National Institutes of Health Guide for the Care and Use of Laboratory Animals. For initial metabolic cage experiments, we used adult (8-12-week-old) C57BL/6J mice (The Jackson Laboratory, Stock 000664). For natural, chemogenetic, and optogenetic torpor experiments, we used 8-12-week-old *Vglut2-IRES-Cre* (The Jackson Laboratory, Stock 028863), *Adcyap1-2A-Cre* (The Jackson Laboratory, Stock 030155), and *Lepr-IRES-Cre* (The Jackson Laboratory, Strain 008320) mice. For adrenergic necessity experiments, we utilized 8-12-week-old β1/β2-KO animals (The Jackson Laboratory, Stock 003810). All mice were housed at 22°C under a standard 12 h light/dark cycle until they were subjected to surgery, at which point they were housed under a reversed 12 h light/dark cycle. Power analysis was used to predetermine sample size, where appropriate (e.g radioactive glucose uptake as well as hyperinsulinemic-euglycemic clamp experiments). Mice were randomly assigned to experimental groups before surgery. Where possible, investigators were blinded during analysis.

### Telemetric monitoring of core body temperature and gross motor activity

Mice were implanted abdominally with telemetric temperature and activity probes (Unified Information Devices, UCT-2112). After at least four days of recovery, animals were singly housed and recorded in standard cages placed onto a radiofrequency receiver platform (Unified Information Devices, UID Mouse Matrix). Core body temperature and gross motor activity were logged every 300 ms based on animal movement and averaged over either 1- or 5-minute windows, as appropriate. Temperature data were collated and exported from the MouseMatrix Software (Unified Information Devices, MouseMatrix 1.6.2) for downstream processing. Analysis of temperature data was performed in R (R Core Team) using a custom-written R Script.

### Metabolic cage experiments

Metabolic cage data was collected on individually housed mice placed in a Promethion indirect calorimeter (Sable Systems) with a temperature-controlled cabinet (Pol-Eco) and provided with ad libitum food (Labdiet 5008) and water purified by reverse osmosis. Starr Scientific telemetry receiver bases were placed beneath the Promethion Cages to match the implanted probes. Mice were maintained under 12 h light/dark photoperiods (0600–1800) at an ambient temperature of 23 ± 0.2°C. Position and physical activity were collected every second. Rates of oxygen consumption (VO_2_) and carbon dioxide production (VCO_2_) were measured every 2 min. Energy expenditure was calculated with the Weir equation^40^.

### Fasting-inducted torpor

Telemetric temperature probe-implanted adult (8-18-week-old) mice were singly housed before the induction of torpor. Each mouse was moved to a new individual cage containing water and nesting material but devoid of bedding and food at the beginning of the dark cycle (ZT 11.5). Initial bouts of torpor were observed after approximately 8-12 h of fasting. Mice were returned to their standard cages containing food 24 h after the start of the fast. The ambient temperature of the facility was maintained at ∼22°C.

### Stereotactic viral injection and fiber implantation

For injections, mice were anaesthetized with 3% isoflurane and placed in a stereotaxic head frame (Kopf Instrument, model 1900). Animals were kept constantly sedated using 1-1.5% isoflurane, and either Buprenorphine SR (1.0 mg/kg) or Ethiqa SR (3.25 mg/kg) were administered for analgesia. The skull was leveled in the AP and ML directions, and bore holes were drilled using a stereotaxic drill (David Kopf Instruments). For chemogenetic experiments, unless otherwise specified, craniotomies and viral administration were bilateral. For optogenetics, craniotomies and viral administration were unilateral. An air-based injection system built with Digital Manometer (Grainger, 9LHH8) was used to infuse the virus through pulled glass capillaries. Virus was administered at the following coordinates - POA: AP +0.4 mm, ML ±0.5 mm, DV −5.02 mm, PVH: AP -1.0 mm, ML ±0.1 mm, DV −4.5mm, DMH: AP - 1.8 mm, ML ±0.5 mm, DV −5.02 mm, PAG: AP -3.25 mm, ML ±0..25 mm, DV −1.8 mm, RPa: AP -5.5 mm, ML ±0.0 mm, DV −5.75 mm. Virus was infused at approximately 100 nL min^−1^, and the needle was kept at the injection site for 10 min before withdrawal. Unless otherwise specified, 100 nL of virus was administered per site (unilateral: 100 nL total, bilateral 200 nL total). Following needle removal, the incision was closed using 4-0 monocryl adsorbable suture (Ethicon) with alternating box knots in an interrupted pattern.

For optic fiber implantation for optogenetics, 400 μm, 0.37 NA borosilicate mono fiber optic cannulae (Doric, MFC_400/430-0.37_###_MF1.25_FLT, where ### corresponds to fiber length) were implanted approximately 500 μm above the injection site. Fibers were initially fixed to the head using superglue (Gorilla Super Glue Gel), followed by Metabond (CB Metabond S380) to ensure the stability of the implant. For optogenetics experiments, 100 nL of a 1:1 mixture of AAV8-hSyn-DIO-hM3D(Gq)–mCherry, or AAV8-hSyn-DIO-hM3D(Gq)–mCherry and AAV8-EF1a-ChR2(H134R)-EYFP, respectively, were injected unilaterally into the POA. Chemogenetic stimulation of POA neurons followed by monitoring of core temperature decreases was used to validate injection accuracy. Animals not demonstrating temperature reductions upon administration of CNO were removed from the analysis.

### Viral constructs

AAV8-hSyn-DIO-hM3D(Gq)-mCherry (Addgene, 44361-AAV8), AAV-hSyn-fDIO-hM3D(Gq)-mCherry-WPREpA (Addgene, 154868-AAV8), pAAV-EF1a-double floxed (DIO)-hChR2(H134R)-EYFP-WPRE-HGHpA (Addgene, 20298-AAV1), pAAV-hSyn-DIO-hM4D(Gi)-mCherry (Addgene, 44362-AAV8), AAV-CaMKIIa-hM3D(Gq)-mCherry (Addgene, 50476-AAV8), AAV-hSyn-hM3D(Gq)-mCherry (Addgene, 50474-AAV8), AAV-hSyn-DIO-mCherry (Addgene, 50459-AAV8), and pAAV-hSyn-DIO-EGFP (Addgene, 50457-AAV8) were obtained from Addgene. AAV-EF1a-DIO-EYFP (AAV2) was prepared by the UNC Viral Core. AAV-DIO-TK-P2A-EGFP and HSV-H129-ΔTK-tdTomato were prepared by the University of California Irvine Viral Core.

All AAV viruses were diluted with PBS to a final concentration between 5 × 10^12^ and 1 × 10^13^ genome copies per mL before stereotactic delivery into the mouse brain. HSV-H129-ΔTK-tdTomato was administered at 5 × 10^8^ PFU per mL (100 nL unilaterally).

### Chemogenetic induction and inhibition of torpor

#### Induction

Cre-expressing animals were bilaterally transduced with AAV8-hSyn-DIO-hM3D(Gq)-mCherry. Gq-DREADD-transduced animals were implanted with telemetric temperature probes and allowed to recover for a minimum of 5 days post-surgery. Following this recovery window, animals were injected with CNO, and body temperature was monitored to confirm accuracy of injection. Animals were considered to be torpid if their body temperatures decreased below 34°C upon CNO administration.

CNO solution was prepared by initially dissolving CNO hydrochloride (Sigma-Aldrich, SML2304) in H_2_O to generate a stock solution of 5 mg/mL. The stock solution was diluted with PBS to a final concentration of 0.2 mg/mL, and approximately 125 µL was injected intraperitoneally per mouse for a final injection concentration of 1 mg kg^−1^ in bilaterally injected animals. Unilaterally injected animals received concentrations of 2 mg kg^-1^.

#### Inhibition

Cre-expressing animals were bilaterally transduced with AAV8-hSyn-DIO-hM4D(Gi)-mCherry. Gi-DREADD-transduced animals were implanted with telemetric temperature probes and allowed to recover for a minimum of 5 days post-surgery. Following this recovery window, animals were first fasted overnight starting in the dark phase (ZT12-12). Animals were injected with PBS at ZT22, right before the onset of the light phase, and body temperature was monitored to confirm accuracy of injection. Animals were considered to be torpid if their body temperatures decreased below 34°C. Animals were recovered for 7 days, and subjected to this fasting paradigm again, except CNO (10 mg/kg) was injected at ZT22. Injections were considered successful if animals entered torpor when injected with PBS, but did not when injected with CNO. If entry into torpor was not inhibited by CNO the animals were excluded from further experiments.

### Optogenetic stimulation of avPOA^Vglut2^ neurons

Animals expressing either ChR2(H134R) or an EYFP control, and implanted with a fiber optic cannula, were allowed to recover 5 days post-surgery before confirming proper injection location via CNO stimulation. Animals that failed to demonstrate a body temperature decrease were excluded from further analysis. After a 2-week post-surgery recovery period to allow for adequate viral expression, awake, freely moving animals were tethered to a fiber optic patch cord (Doric) and placed in a deep-well cage (WPI) with free access to food and water. Animals were allowed to acclimate to the new environment for 1 h prior to the start of stimulation. To deliver light pulses a PulsePal2 (SanWorks) was utilized as a pulse train generator, which was connected to a desktop computer via USB and controlled via a custom Python script. Pulses from the controller were conveyed via a 455 nm fiber-coupled LED (ThorLabs M455F3) connected to the mouse via patch cords connected through a rotary joint (ThorLabs RJ1) to allow for unencumbered movement of the animal. For ChR2 experiments, the stimulation paradigm was as follows: 8.5 mW, 10 µs pulse, 50 µs interval (16 Hz), Duty Cycle 1s ON 1s OFF. Power output was confirmed at the fiber tip via a power meter (ThorLabs PM100D) prior to each experiment. For fasting experiments, such as GTT or overnight fasting, an identical setup was utilized, but the animal was not given food (access to water was maintained).

Following the completion of optogenetic experiments, animals were sacrificed, perfused with 4% PFA, and their brains were sectioned to confirm the accuracy of fiber placement.

### Intraperitoneal glucose tolerance tests (GTT)

For chemogenetic glucose tolerance testing, Gq-DREADD or Gi-DREADD-expressing and control animals were weighed and moved to a clean cage containing a bedding square and water, but no food, prior to the beginning of the dark cycle (ZT 11.5), and placed onto telemetric temperature platforms. After 3 hours of fasting and a “Pre-CNO” blood glucose reading (Bayer Onetouch), animals were injected with 1 mg kg^-1^ CNO and returned to their cage for 60 minutes. A baseline blood glucose reading was subsequently taken (t = 0), and animals were injected with 2 g/kg glucose (10*Body Weight 20% glucose). Blood glucose readings were then taken at 15, 30, 45, 60, 90, and 120 minutes post-glucose injection prior to returning animals to their home cages.

For optogenetic glucose tolerance tests, ChR2(H134R)- or EYFP-expressing animals were weighed and connected to a fiber optic patch cord and placed in a clean cage with a nesting square and water but no food. Animals were fasted for 3 hours, sampled for blood, and injected with a volumetrically equivalent amount of saline to CNO that would have been injected in a chemogenetic GTT (i.e. 5 µL saline/g of mouse). After a 60-minute interval, a baseline blood sample was taken, and animals were injected with 2 g/kg glucose. Immediately upon glucose injection, blue light (455 nm) stimulation commenced. Blue light stimulation was continued throughout the duration of the experiment, and blood was sampled at the time points previously described. Blue light stimulation was terminated after t = 120 minutes, and animals were disconnected and returned to their home cages.

Data were recorded, graphed, and analyzed in GraphPad Prism. AUC calculations were performed using GraphPad Prism analytical tools.

### Radioactive glucose uptake

#### Radioactive glucose uptake assay

Male 8-12-week-old *Vglut2-IRES-Cre* animals were transduced with either mCherry or Gq-DREADD. After validating Gq-DREADD animals’ ability to undergo chemogenetic topor, animals were weighed, transferred to a new cage, and fasted for 3 hours. Animals were administered CNO and allowed to equilibrate body temperature for 60 minutes, upon which time initial blood glucose and scintillation counting samples were taken and animals were injected with a mixture of 20 μCi 2-[1,2-^3^H(N)]-Deoxy-D-glucose (^3^H-2-DG), 0.5 U/kg Insulin, and 1 mM 2-Deoxy-Glucose as per Møller et al^41^. Animals were then placed in a new cage, and 3 μL of blood were taken for scintillation counting along with 1 drop for blood glucose measurements at 5, 10, 20, and 30 minutes post-injection. Blood collected for scintillation counting was added to 5 mL of Opti-Fluor (Perkin-Elmer), mixed, and placed into a Perkin-Elmer Imager for scintillation counting. Following 30-40 minutes of uptake, animals were euthanized, and their gastrocnemius, soleus, and BAT were dissected and snap-frozen in liquid nitrogen. Scintillation samples were measured for tritium over 1 minute, and plasma tritium concentration were recorded as disintegrations per minute (DPM).

#### Determination of tissue-specific glucose uptake rate

To determine tissue-specific glucose uptake rate, we performed chromatographic separation of ^3^H-2-DG from ^3^H-2-DG-6-Phosphate, based on a protocol developed by Kim et al^20^. Using ^3^H-2-DG-6-Phosphate as a proxy for tissue-specific uptake is predicated on the fact that ^3^H-2-DG is metabolized to ^3^H-2-DG-6-Phosphate almost immediately upon entry into the cell, which then precludes its exit from the tissue. Measurement of total ^3^H-2-DG includes interstitial, unmetabolized ^3^H-2-DG, and is therefore susceptible to artefactual over- or under-estimation of glucose uptake based on state-dependent differences in tissue perfusion. To avoid this confound, we utilized Poly-Prep anion exchange columns (Bio-Rad, 731-6211) to isolate the ^3^H-2-DG-6-Phosphate. Briefly, tissues were weighed and homogenized in 10 volumes of sterile, deionized water. Homogenized tissues were then heated to 95°C for 10 minutes, cooled to room temperature, vortexed, and centrifuged for 10 minutes at 21430 RCF. Subsequently, 33.3 μL of homogenate was added to 4.967 mL of Opti-Fluor for scintillation counting. During this time, the Poly-Prep anion exchange column was equilibrated with 6 mL of sterile deionized water. Following equilibration, 333 μL of homogenate was applied to the columns and allowed to flow through by gravity. The column was subsequently washed 3 times with 2 mL of sterile, deionized water, and the flowthrough was collected in scintillation vials as the “wash” fraction. Finally, the ^3^H-2-DG-6-Phosphate was eluted from the column using a 0.3 M Formic Acid/0.5 M NH_4_CH_3_COO elution buffer. Elution was accomplished via 3 washes of 2 mL using the elution buffer and collected in scintillation vials. The wash and elution (500 μL) were added to 5 mL of Opti-Fluor, and the samples were vortexed and analyzed via scintillation counting.

For quantitation of tissue-specific glucose uptake rates, we employed a method modified from Møller et al^41^. Briefly, the DPM of the eluate, representing the ^3^H-2-DG-6-Phosphate concentration in the tissue, was related to the time of uptake and weight of tissue to obtain a DPM g^-1^ min^-1^ value, representing tissue-specific uptake rate. The time-weighted ^3^H-2-DG appearance was then extrapolated from the DPM of ^3^H-2-DG in the blood and divided by the time-weighted total glucose concentration to obtain the specific activity of ^3^H-2-DG in the blood (DPM/μmol). The glucose uptake index for the tissue was then calculated by dividing the tissue uptake rate by the specific activity, and expressing this value as nmol g^-1^ min^-1^.

### Immunoblotting

Male, 8-12-week-old *Vglut2-IRES-Cre* animals validated to have body temperature drops and glucose intolerance when stimulated with CNO were subdivided into three cohorts, injected with either saline, CNO (1 mg/kg), or insulin (0.5 IU/kg), respectively, and monitored for blood glucose changes over a 45-minute period. After 45 minutes, animals were sacrificed, and gastrocnemius, soleus, quadriceps, liver, and BAT were dissected and snap-frozen. These tissues were then homogenized in Radioimmunoprecipitation (RIPA) Buffer composed of 150 mM NaCl, 1.0% (w/v) Triton X-100, 0.5% (w/v) Sodium Deoxycholate, 0.1% (w/v) sodium dodecylsulfate, 50 mM Tris (pH 8), EDTA-free protease inhibitor (Roche), and PhosSTOP phosphatase inhibitor (Roche), using a BioRuptor (Qiagen). Lysates were cleared by centrifugation in a microcentrifuge (21430 RCF for 10 min at 4°C), and protein concentrations were quantified using the Pierce BCA Protein Assay Kit (Thermo Fisher). Lysate samples were prepared by the addition of 5× Laemelli Buffer (0.242 M Tris, 10% SDS, 25% glycerol, 0.5 M dithiothreitol (DTT) and bromophenol blue), resolved by Bis-Tris SDS–PAGE, and analyzed by immunoblotting. Antibodies and concentrations used for immunoblotting include Insulin Receptor β (4B8) Rabbit mAb #3025 (Cell Signaling Technologies) 1:1000, Akt Antibody #9272 (Cell Signaling Technologies) 1:1000, and Phospho-Akt (Ser473) Antibody #9271 (Cell Signaling Technologies) 1:500. Proteins were visualized using an Anti-rabbit IgG HRP-linked Antibody #7074 (Cell Signaling Technologies) 1:3000 in combination with the chemiluminescent Pierce™ ECL Western Blotting Substrate (ThermoFisher) using an X-OMAT film processor.

### Sciatic nerve denervation

Sciatic nerve denervation procedures were modeled on those described in Volodin et al^42^. Briefly, animals were anesthetized with 3% isoflurane and administered analgesics. The sciatic nerve was then accessed via the back right flank of the animal and severed at both distal and proximal ends, such that there was no reattachment of nerve fibers during the course of recovery. Accuracy of the cut was validated via observation of the animal post-operation. Severing the sciatic nerve causes a stereotyped dragging of the ipsilateral limb. Animals confirmed to have successful denervations were included in experimental cohorts, whereas those that retained limb mobility were excluded. Animals were allowed to recover for 72 hours prior to use in radioactive glucose uptake assays previously described.

When dissecting organs for analysis of tissue-specific glucose uptake, both innervated and denervated gastrocnemius muscles were harvested. These tissues were processed in an identical manner, and the ratio of innervated:denervated glucose uptake was used as a metric for the reduction in glucose uptake in the same animal that resulted from denervation.

### Blood collection

Tail blood was collected from animals by opening a small hole in the tail of the animal and manually extracting 20 μL of blood into a Lithium Heparin Microvette (Sarsdedt). For longitudinal assays, no more than 150 μL of blood was removed from an animal. Following blood extraction, microvettes were placed on ice for no longer than 60 minutes, and then centrifuged in a microcentrifuge for 5 minutes at 2000 RCF to separate blood cells from plasma. Plasma was extracted from these tubes, aliquoted, snap-frozen in liquid N_2_, and placed at -80°C for long-term storage.

### ELISA assays

All assays were performed according to manufacturers’ instructions, as briefly described below:

#### Insulin

Following plasma separation, 10 μL plasma from each mouse was used for the Mouse Ultra-Sensitive ELISA Assay (Crystal Chem), plated in 5 μL duplicates. Samples were compared against a standard curve and analyzed using a Spectramax iD3 plate reader with a reference wavelength of 630 nm and a measured wavelength of 450 nm. Data were fit using a linear curve, and cohort analysis was performed using two-tailed t-tests.

#### Glucagon

Following plasma separation, 20 μL plasma from each mouse was used for the Mouse Glucagon ELISA Assay (Crystal Chem), plated in 10 μL duplicates. Samples were compared against a standard curve and analyzed using a Spectramax iD3 plate reader with a reference wavelength of 630 nm and a measured wavelength of 450 nm. Data were fit using a 4-parameter logistic curve, and cohort analysis was performed using two-tailed t-tests.

#### Leptin

Following plasma separation, 10 μL plasma from each mouse was used for the Mouse Leptin ELISA Assay (Crystal Chem), plated in 5 μL duplicates. Samples were compared against a standard curve and analyzed using a Spectramax iD3 plate reader with a reference wavelength of 630 nm and a measured wavelength of 450 nm. Data were fit using a 4-parameter logistic curve, and cohort analysis was performed using two-tailed t-tests.

#### Corticosterone

Following plasma separation, 5 μL plasma from each mouse was used for the Corticosterone ELISA Assay (Enzo Life Sciences). Samples were enriched for steroids using the provided Steroid Displacement Reagent and plated in duplicate. Samples were compared against a standard curve and analyzed using a Spectramax iD3 plate reader with a reference wavelength of 580 nm and a measured wavelength of 405 nm. Data were fit using a 4-parameter logistic curve, and cohort analysis was performed using two-tailed t-tests.

### Polar serum metabolomics

To begin, 10 μL aliquots of plasma from blood samples was combined with 90 μL of 75:25:0.2 acetonitrile:methanol:formic acid containing 0.2 μg/mL internal standards and vortexed for 5 minutes as 4°C. Subsequently, the samples were spun at 21430 RCF for 10 minutes in a refrigerated centrifuge at 4°C. Supernatant was taken, and 15 μL was added to LC/MS vials for analysis.

Analysis was conducted on a QExactive benchtop Orbitrap mass spectrometer equipped with an Ion Max source and a HESI II probe, which was coupled to a Dionex UltiMate 3,000 UPLC system (Thermo Fisher Scientific). External mass calibration was performed using the standard calibration mixture every 7 days. Chromatographic separation was achieved using the following conditions: buffer A was 20 mM ammonium carbonate, 0.1% ammonium hydroxide; buffer B was acetonitrile. The column oven and autosampler tray were held at 25°C and 4°C, respectively. The chromatographic gradient was run at a flow rate of 0.150 mL min−1 as follows: 0–20 min: linear gradient from 80% to 20% B; 20–20.5 min: linear gradient from 20% to 80% B; 20.5–28 min: hold at 80% B. The mass spectrometer was operated in full-scan, polarity switching mode with the spray voltage set to 3.0 kV, the heated capillary held at 275°C, and the HESI probe held at 350°C. The sheath gas flow was set to 40 units, the auxiliary gas flow was set to 15 units, and the sweep gas flow was set to 1 unit. The data acquisition was performed over a range of 70–1,000 m/z, with the resolution set at 70,000, the automatic gain control target at 10E6, and the maximum injection time at 20 ms. Relative quantitation of polar metabolites was performed with TraceFinder 2.2 (Thermo Fisher Scientific) using a 5 p.p.m. mass tolerance and referencing an in-house library of chemical standards.

### Hyperinsulinemic-euglycemic clamp

Survival surgery, to place an indwelling intravenous catheter, was performed 5∼6 days prior to the insulin clamp experiments^20^. On the day of the euglycemic clamp experiment, mice were placed in a rat restrainer with their tails tape-tethered at one end to minimize animal stress and allow access to tail blood sampling during the experiment. Mice were allowed to acclimatize to this state for 2 hours prior to the insulin infusion. During this 2-h acclimation period, D-[3-^3^H] glucose was infused (0.05 µCi/min) using a microdialysis pump to assess the basal rate of whole-body glucose turnover (i.e, basal hepatic glucose production). Subsequently, a baseline blood sample (60 µL) was collected for the measurement of basal plasma glucose, insulin, and [^3^H] glucose concentrations^20^.

Following the basal period, a 2-h hyperinsulinemic-euglycemic clamp was conducted with a primed (150 mU/kg body weight) and continuous infusion (2.5 mU/kg/min) of human insulin to raise plasma insulin within a physiological range (approximately 300 pM)^20^. Blood samples (10 μL) were collected at 10∼20-minute intervals for the measurement of plasma glucose levels. Based on plasma glucose levels, 20% glucose was infused at variable rates to maintain euglycemia (∼120 mg/dL) throughout the clamp. To determine insulin-stimulated whole-body glucose turnover, continuous infusion of [3-^3^H] glucose was applied throughout the clamp (0.1 µCi/min). To estimate insulin-stimulated glucose uptake in individual organs, a bolus of 2-deoxy-d-[1-^14^C] glucose (2-[^14^C]DG) (10 µCi) was administered at 75 min after the start of the clamp^20^. Blood samples (10 µL) were taken at 80, 85, 90, 100, 110, and 120 min for the measurement of plasma [^3^H] glucose, ^3^H_2_O, and 2-[^14^C] DG concentrations. A final blood sample (20 µL) was taken at the end of the clamp to measure plasma insulin concentrations. At the end of the insulin clamp, mice were anesthetized, and skeletal muscles (gastrocnemius and quadriceps) from both hindlimbs, white and brown adipose tissues, liver, brain, and heart were collected for biochemical analysis^20^. The tissues were rapidly frozen in liquid N_2_, and stored at −80°C until biochemical analysis was performed. Tissue-specific uptake was determined as described in **Radioactive glucose uptake** above.

### Phosphoproteomics

#### Proteolytic digestion and peptide extraction

Tissues were lysed at 70°C for 15 min in 1 mL 1X SDS lysis buffer (5 % (w/v) SDS, 100 mM TEAB pH 8.5, 40 mM CAA, 10 mM TCEP, protease inhibitor cocktail, phosphatase inhibitor cocktail) in Low Protein Binding Microcentrifuge Tubes (Thermo Scientific, Waltham, MA, USA). Cellular debris was removed by centrifugation at 15,000 x g for 5 min. Protein (600 µg) was purified via the SP3 method as in Hughes et al^43^. Proteins were bound to 120 µL SP3 beads by adding 2.4 mL 100 % (v/v) ethanol followed by three washing steps with 1 mL 80 % (v/v) ethanol. Samples were digested with trypsin/LysC mix (1:50) in 300 µL 100 mM TEAB, and proteins were digested overnight in a shaking incubator at 37°C at 115 RPM. The following day, an additional 6 µg trypsin/Lys-C mix was added in 6 µL 100 mM TEAB and the digestion continued for 4 hours at 37°C. Peptide digests (240 µg) were dried in a speed-vac concentrator and resuspended in 40 µL 100 mM TEAB. Peptides were labeled with 500 µg TMTpro label resuspended in 20 µL anhydrous acetonitrile. TMTpro-labeled samples were quenched for 30 min at room temperature with 5 µL 5% hydroxylamine in 100 mM TEAB. Labeled peptides were pooled together and desalted on a 500 mg C18 SepPak column (Waters, Milford, MA, USA).

The TMTpro pooled sample was split, with one part used for proteome analysis and the other part used for phosphopeptide enrichment. The proteome sample was fractionated using a Strong Anion Exchange column (Thermo Scientific, Waltham, MA, USA). Increasing concentrations of ammonium acetate (0, 20, 50, 100, 200, 500 mM) were used for elution. Low salt fractions (0, 20, 50 mM ammonium acetate) and high salt fractions (100, 200, 500 mM ammonium acetate) were pooled, respectively, and lyophilized. The phosphopeptide sample was enriched using the High Select™ TiO_2_ phosphopeptide enrichment kit according to the manufacturer’s instructions (Thermo Scientific, Waltham, MA, USA). All TiO_2_ flowthrough and wash fractions were pooled, lyophilized, and further enriched using the High Select™ Fe-NTA phosphopeptide enrichment kit according to the manufacturer’s instructions (Thermo Scientific, Waltham, MA, USA). The low salt and high salt proteome samples, as well as both phosphopeptide-enriched samples, were then fractionated with the High pH Reversed-Phase Peptide Fractionation Kit (Thermo Scientific, Waltham, MA, USA) using the following 12-step gradient of increasing acetonitrile concentrations: 5, 7.5, 10, 12.5, 15, 17.5, 20, 22.5, 25, 27.5, 30, 60%. The following fractions were then pooled together and lyophilized: 1+7, 2+8, 3+9, 4+10, 5+11, 6+12. Lyophilized fractions were resuspended in 12 µL 0.2% formic acid in MS-grade water for LC-MS analysis.

#### LC-MS data collection

Mass spectrometry was performed using an Orbitrap Eclipse mass spectrometer equipped with a FAIMS Pro source, connected to a Vanquish Neo nLC chromatography system using an EasySpray ES902 column (75 µm x 25 cm, 100 Å), all from Thermo Fisher Scientific (Waltham, MA, USA). Peptides were separated at 300 nL/min on a gradient of 3–25% B for 90 min, 25–40% B for 30 min, 40–95% B for 10 min, 95% B over 6 min, using 0.1% FA in water for A and 0.1% FA in 80% acetonitrile for B. The Orbitrap and FAIMSpro were operated in positive ion mode with a positive ion voltage of 2100 V, an ion transfer tube temperature of 305°C, and a 4.2 L/min carrier gas flow, using standard FAIMS resolution and compensation voltages of -45, –55, and –65 V. Full scan spectra were acquired in profile mode at a resolution of 120,000 (MS1 and MS2), with a scan range of 400–1400 m/z, custom maximum fill time (200 ms), custom AGC target (100% MS1, 250% MS2), isolation windows of m/z 0.7, intensity threshold of 2.0e4, 2–6 charge state, dynamic exclusion of 60 seconds, and 35% HCD collision energy.

#### Proteomic and Phosphoproteomic Data Analysis

Proteome data were analyzed using the PEAKS Studio 10.6 software package. Raw data extracted from .raw files were pre-processed with the following settings: scans were merged within a 10 ppm retention time window and a 10 ppm precursor m/z tolerance, including precursor mass and charge states (z = 2-8). Pre-processing steps included automatic centroiding, deisotoping, and deconvolution.

Protein identification was performed using a mouse protein FASTA database (Uniprot Taxon ID 10090), with the following parameters: parent mass error tolerance set to 10 ppm, fragment mass error tolerance set to 0.05 Da, retention time shift tolerance of 5.0 minutes, and semispecific trypsin enzyme specificity. TMT was applied as an unvariable modification, while the variable modifications considered included carbamidomethylation (C), oxidation (M), phosphorylation (P), and deamidation (NQ). Non-specific cleavage at one terminus was allowed, with up to three missed cleavages and a maximum of three variable posttranslational modifications per peptide. The false discovery rate (FDR) was estimated using the decoy-fusion approach, with peptides having an FDR ≤ 1% and a significance threshold of 20 (-10lgP) considered confidently identified. For protein quantification, at least two high abundant peptide signals were selected to calculate raw protein peak areas. The software computed phosphosite data by using the abundances of multiple peptides for each phosphorylation.

## Acknowledgements

We thank T. Diefenbach for his help with whole brain imaging experiments. We thank all Hrvatin Lab members for their insightful comments on the manuscript. Whole brain imaging was performed at the Ragon Institute Microscopy Core. Metabolomics experiments were performed with the Whitehead Institute Metabolomics Core. Proteomic and phosphoproteomics experiments were performed with the Whitehead Institute Quantitative Proteomics Core. Metabolic cage experiments were performed at the BIDMC Energy Balance Core. Metabolic clamp experiments were performed at the UMass Metabolic Disease Research Center.

## Funding

Mathers Foundation Grant (S.H.)

Searle Scholars Program (S.H.)

Pew Charitable Trust (S.H.)

Rosenkranz Foundation (S.H.)

MIT Research Support Committee Grant (S.H.)

Howard Hughes Medical Institute Hrabowski Scholar (S.H.)

NIH Director New Innovator Award DP2 DP2DK136123 (S.H.)

## Author Contributions

J.M.R and S.H conceptualized the project. J.M.R, S.H., E.C.G, J.K.K, F.S, and A.S.B devised experimental methodologies. J.M.R, M.A, N.N, H.W, M.M, M.W, B.L, B-Y.K, and B.K executed experiments. J.M.R, S.H, M.A, N.N, and H.W contributed to data visualization. S.H and E.C.G acquired funding. S.H and E.C.G administered and supervised the project. J.M.R, S.H, and E.C.G wrote the initial manuscript draft. J.M.R, S.H, E.C.G, M.A, N.N, H.W, A.S.B, and J.K.K reviewed and edited the manuscript.

## Data and Material Availability

All data needed to evaluate conclusions made in both main text and supplementary figures in this paper will be made available upon reasonable request.

## Supplementary Materials

Materials and Methods

Figs S1 to S7

## Supplementary Figures

**Supplementary Figure 1:**
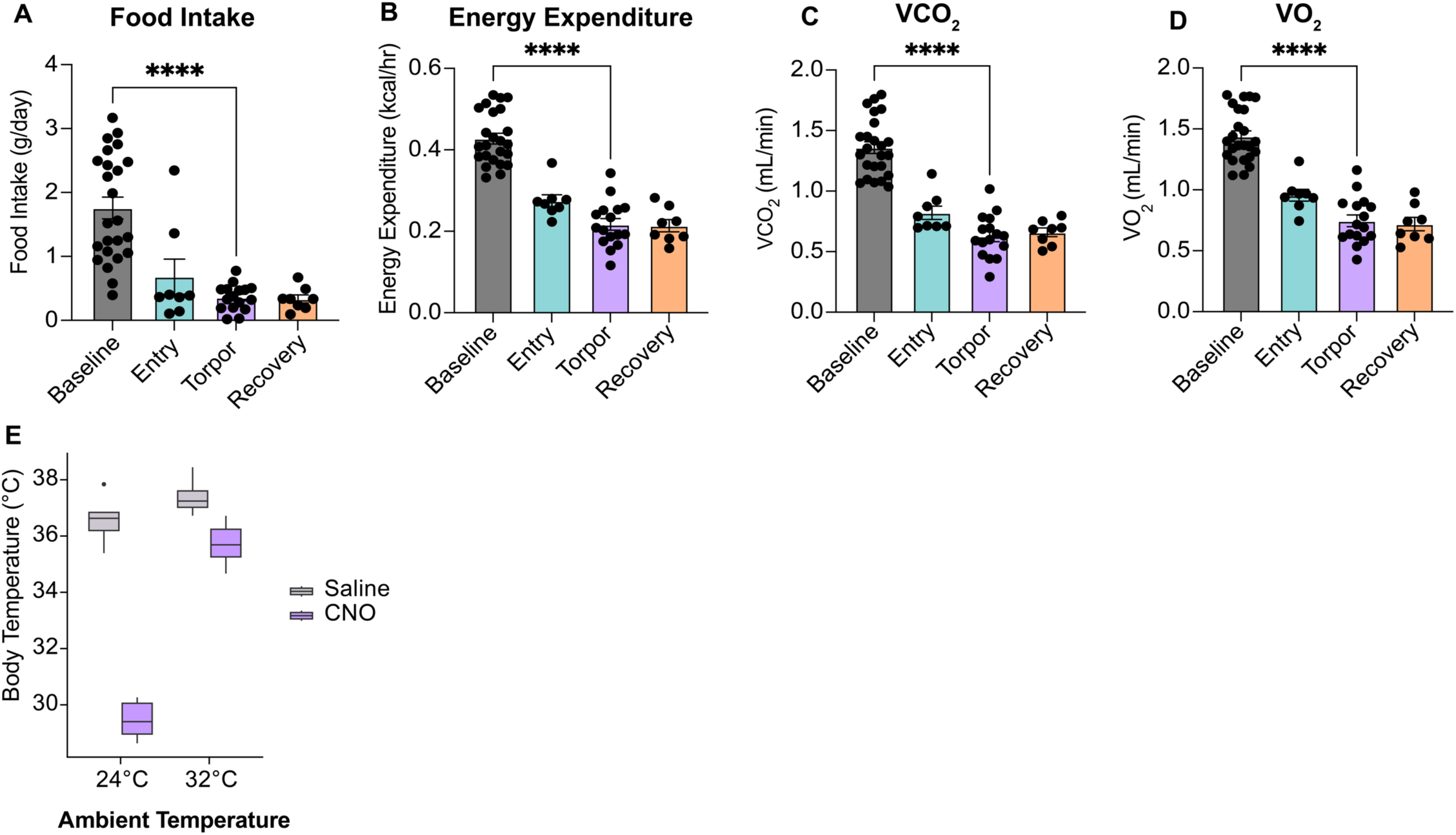
avPOA stimulation regulates diverse metabolic parameters. **(A-D)** Quantitation of (**A)** Food Intake, **(B)** Energy Expenditure, **(C)** VCO_2_, and (**D)** VO_2_ throughout different phases of avPOA stimulation described in Figures 1c-f. Baseline = pre-CNO stimulation, Entry = 60 minutes post-CNO stimulation, Torpor = duration of torpor bout, Recovery = post-torpor period before returning to euthermia (**** = p < 0.00005, mean ±SEM, student’s two-tailed t-test, n = 8 WT C57Bl/6J mice). **(E)** Quantitation of animal body temperature following 300 minutes of saline or Gq-DREADD stimulation at either T_a_ *=*24°C or 32°C (n = 8, mean ±SEM).

**Supplementary Figure 2:**
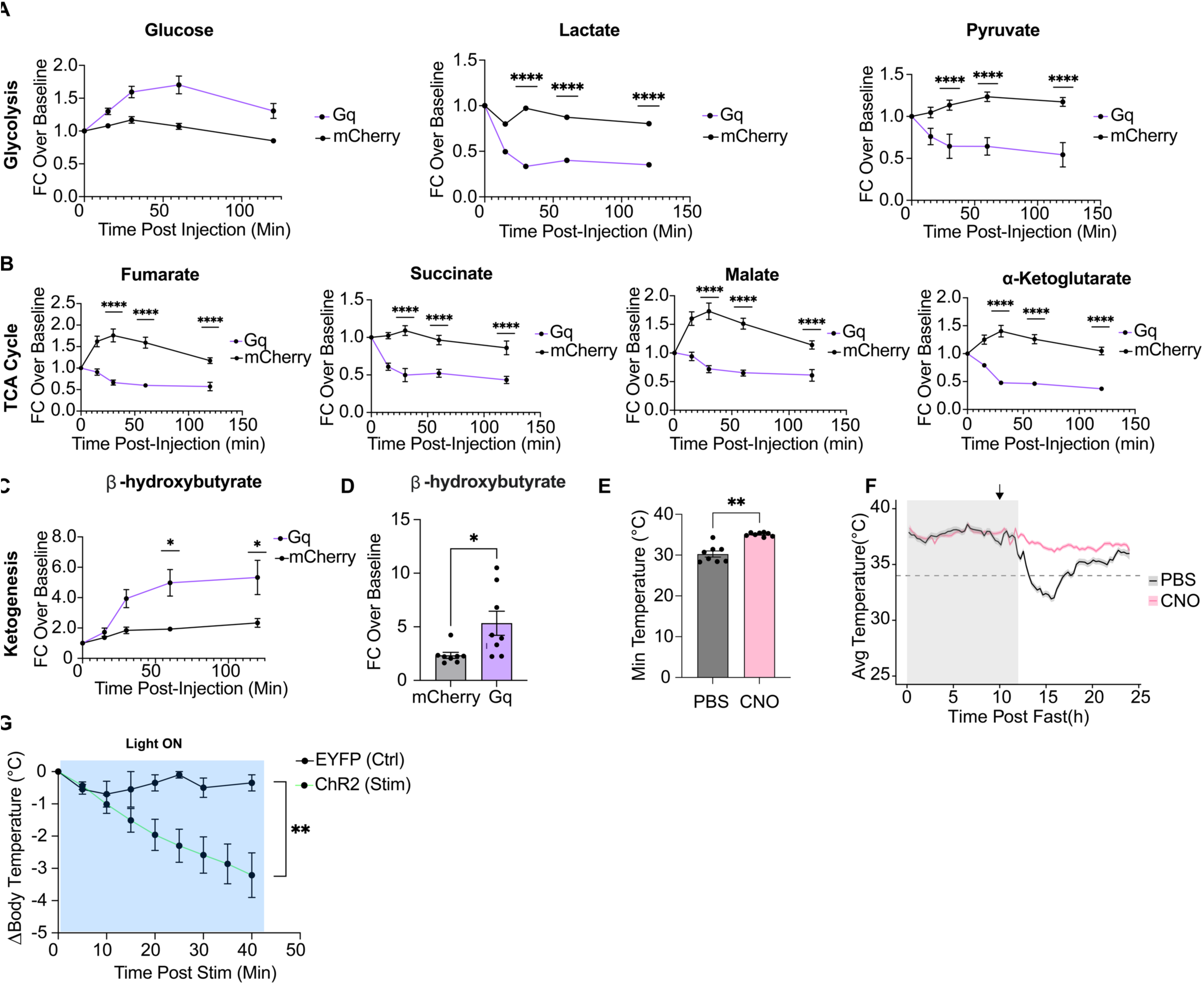
TCA cycle and glycolytic metabolite abundance is altered following CNO stimulation. **(A)** Major glycolytic metabolites, including pyruvate and lactate, are monotonically decreased by CNO stimulation of Cre-dependent mCherry- or Gq-DREADD-transduced *Vglut2-IRES-Cre* mice as compared to their saline-injected counterparts. Conversely, glucose is monotonically increased upon CNO stimulation as compared to saline injection (** = p < 0.005, **** = p < 0.00005, student’s two-tailed t-test, mean ±SEM, n = 8 animals). **(B)** Significantly downregulated TCA cycle metabolites include fumarate, succinate, malate, and α-ketoglutarate (**** = p < 0.00005, student’s two-tailed t-test, mean ±SEM, n = 8 animals). Sample values reported as cohort-specific baseline-normalized fold change. Blood was collected at 0, 15, 30, 60, 120 minutes post-injection. **(C)** β-hydroxybuyrate levels are significantly increased in CNO-injected animals as compared to controls (* = p < 0.05, student’s two-tailed t-test, mean ±SEM, n = 8 animals). **(D**) Quantitation of (C) 120 minutes after CNO stimulation in mCherry and Gq animals (* = p < 0.05, student’s two-tailed t-test, mean ±SEM, n = 8 animals). **(E)** Quantification of minimal T_b_ measured during fasting-induced torpor in Gi-DREADD-transduced *Vglut2-IRES-Cre* animals. Animals were fasted for 24 hours at the beginning of the dark cycle (ZT 12), and either PBS or CNO (10 mg/kg) was injected 10 hours after fasting began (ZT 22) (** = p < 0.005, student’s two tailed t-test, n = 8 PBS, n = 8 CNO, same animals, mean ±SEM). **(F)** Longitudinal trace of animals in (E), demonstrating that Gi-DREADD animals injected with CNO do not go into torpor due to inhibition of avPOA^Vglut2^ neurons, whereas PBS-injected animals enter normally. Arrow corresponds to time of CNO or PBS injection. Gray shading indicates dark cycle (ZT12-24). Horizontal dashed line at 34.5°C corresponds to the temperature threshold for torpor entry. **(G)** Stimulation of avPOA^Vglut2^ neurons with 455 nm light decreases body temperature in ChR2-, but not EYFP-expressing animals over the 40-minute time period of blue light stimulation (* = p < 0.05, student’s two-tailed t-test, mean ±SEM, n = 4 EYFP and n = 5 ChR2 animals).

**Supplementary Figure 3:**
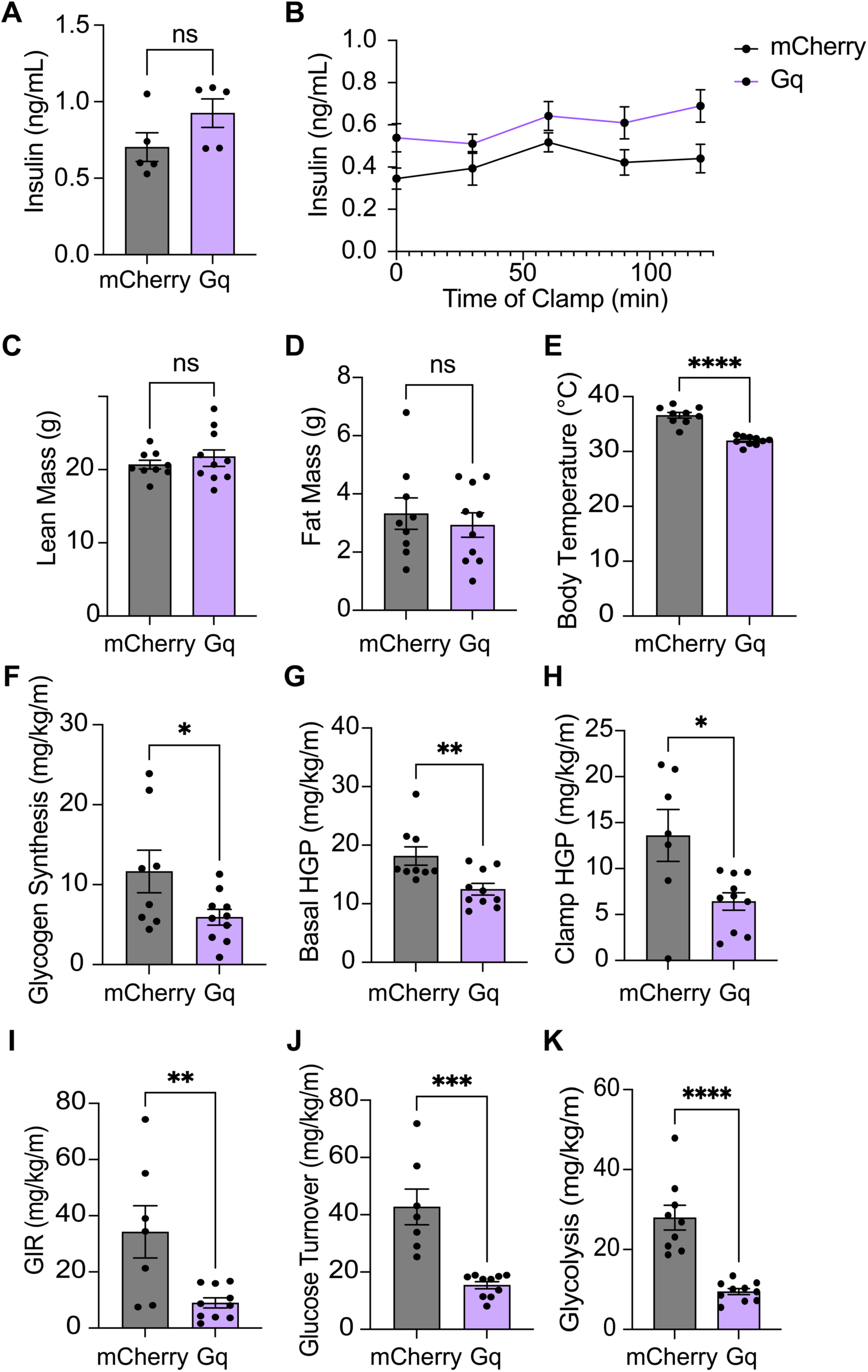
Hyperinsulinemic-euglycemic clamp physical and clamp parameters. **(A)** Quantification of glucose and insulin levels in plasma of fed *Vglut2-IRES-Cre* animals expressing Cre-dependent mCherry or Gq-DREADD in the avPOA, 60 minutes post-CNO injection (p<0.005 = **\*\***). **(B)** Hyperglycemic clamp experiment monitoring plasma insulin levels for 120 minutes during administration of various concentrations of glucose. Blood samples were taken at 0, 30-, 60-, 90-, and 120-minute time points throughout the experiment (n = 9 mCherry, n = 10 Gq-DREADD animals). **(C-K)** correspond to the hyperinsulinemic-euglycemic clamp described in Figure 3. (**C)** Lean mass of animals analyzed by Dual X-ray Absorptiometry (DEXA) prior to beginning of the clamp. (**D)** Fat mass analyzed by DEXA prior to beginning of the clamp. (**E)** Average temperature of animals throughout the duration of the clamp. **(F)** Glycogen synthesis calculated during the clamp based on relative incorporation of ^3^H into hepatic glycogen pool. (**G)** Hepatic Glucose Production (HGP) prior to initiation of the clamp as determined by the ratio of basal ^3^H-Glucose infusion to the specific activity of plasma glucose prior to the initiation of the clamp. (**H)** Insulin-stimulated Hepatic Glucose Production during the clamp, determined by the difference between glucose infusion and whole-body glucose turnover rates. Insulin-stimulated HGP reflects the action of insulin on the liver specifically. **(I)** Glucose infusion rate (GIR) for each cohort. GIR was taken as an average over the duration of the experiment. Decreased GIR in Gq-DREADD-expressing animals indicates insulin insensitivity. Two control animals were excluded from analysis, as GIR measurements were highly irregular. **(J)** Glucose turnover in each cohort throughout the duration of the clamp, calculated as the ratio of ^3^H-glucose infusion to plasma glucose specific activity during the final 30 minutes of the clamp. (**K)** Whole-body glycolysis estimated based on glycogen synthesis, GIR, and glucose clearance rates throughout the experiment (n = 9 mCherry, n = 10 Gq-DREADD, * = p < 0.05, ** = p < 0.005, *** = p < 0.0005, **** = p < 0.00005, two-tailed t-test).

**Supplementary Figure 4:**
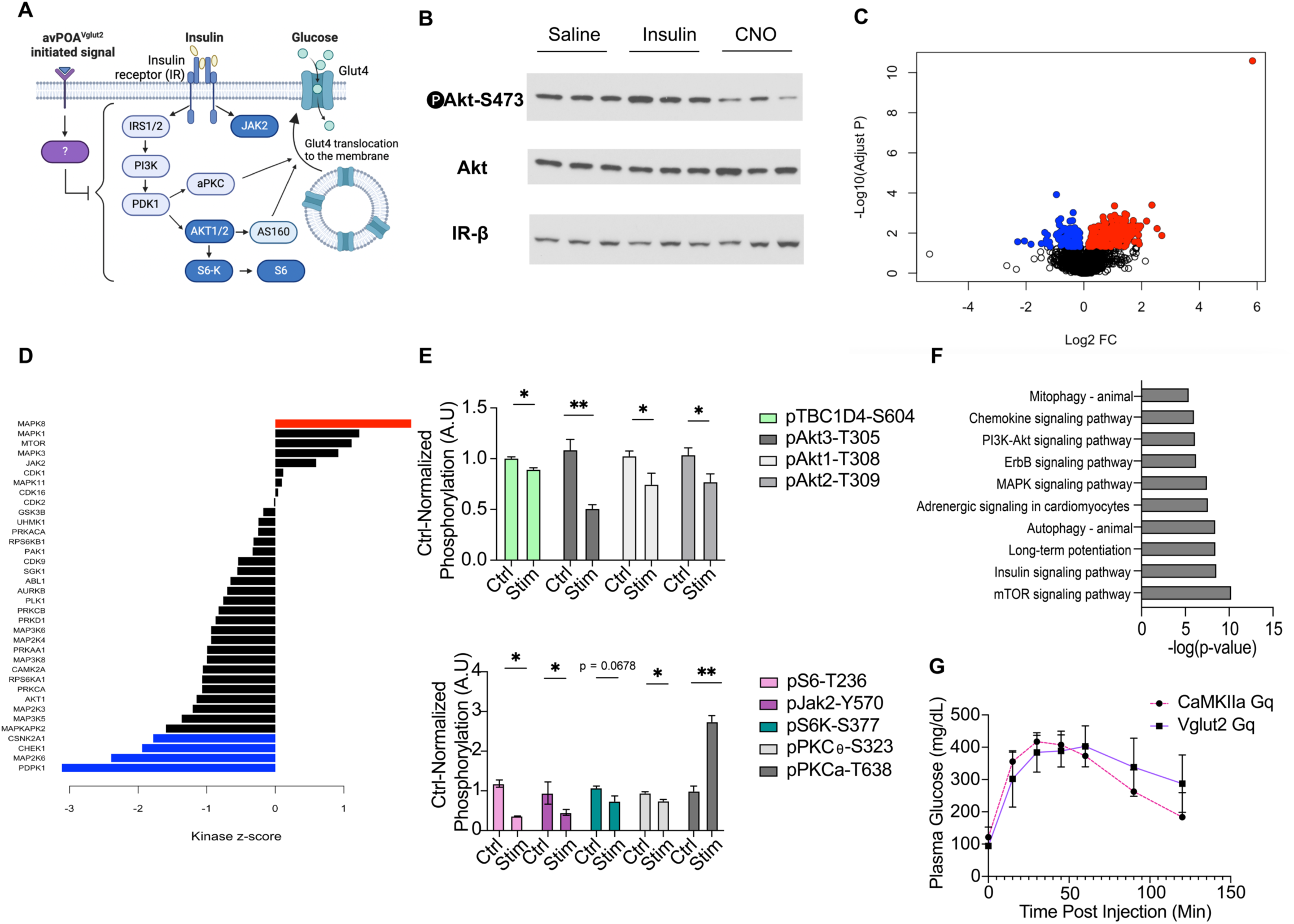
Phosphoproteomic analysis of skeletal muscle reveals avPOA-induced defects in insulin signaling. **(A)** Schematic depicting the insulin signaling pathway and relevant downstream consequences of its activation with respect to glucose uptake. (**B)** Western blot of gastrocnemius muscle lysates obtained from mice injected with either saline, insulin, or CNO showing reduced Akt phosphorylation in CNO-injected animals as compared to saline-injected animals. **(C)** Volcano plot of differentially phosphorylated peptides in the gastrocnemius muscles of *Vglut2-IRES-Cre* mice transduced with Cre-dependent Gq-DREADD in the avPOA and injected with CNO as compared with saline. Significance threshold of p < 0.05 and Log_2_FC of 0.6 used to determine significantly “hyperphosphorylated” or “hypophosphorylated” peptides. Red points correspond to peptides hyperphosphorylated in CNO relative to saline control, while blue correspond to hypophosphorylated peptides. **(D)** Kinase-substrate enrichment analysis score of various kinases identified within the dataset in CNO vs. saline control animals. Kinases whose substrates demonstrated hyperphosphorylation in CNO vs. saline control muscles were scored higher, and those that were significant are shown in red. Kinases whose substrates demonstrated hypophosphorylation in CNO vs. saline control were scored lower, and those that were significant are shown in blue. Z-score cutoff of |1.5| was used to determine significance. **(E)** Saline-normalized phosphorylation of individual residues in key proteins in the insulin signaling pathway (n = 3 animals per cohort, * = p < 0.05, ** = p < 0.005, student’s two-tailed t-test). **(F)** GO-Term analysis of peptides that demonstrated significant (p < 0.05) hypophosphorylation in CNO vs. control animals. **(G)** Comparison of the efficiency of avPOA^Vglut2^ stimulation during a GTT using a CaMKIIa-Gq as compared with a DIO-Gq (AAV-hSyn1-DIO-hM3D(Gq)-mCherry) used in B6 and *Vglut2-IRES-Cre* animals, respectively. No statistically significant difference was observed between the two cohorts.

**Supplementary Figure 5:**
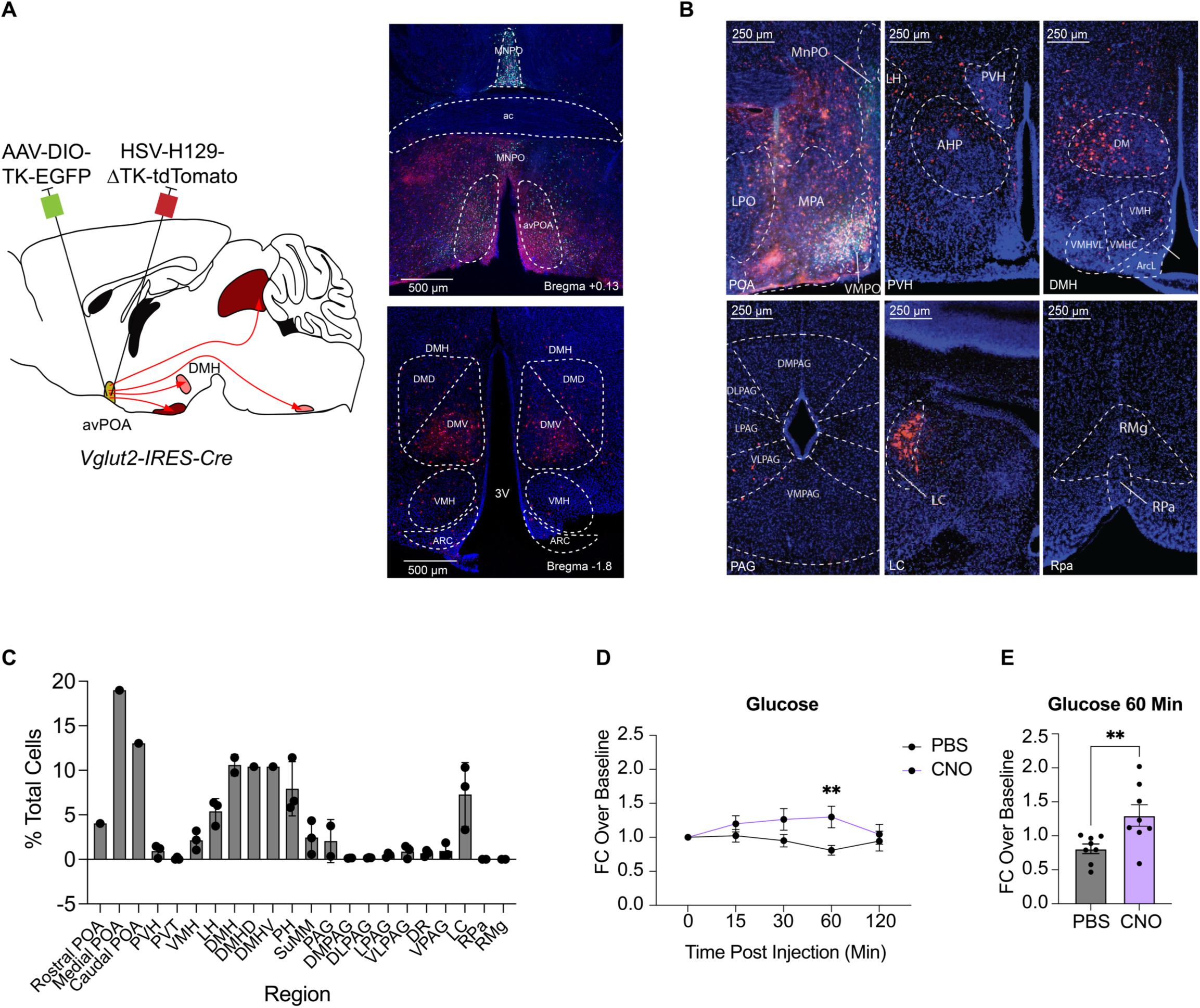
avPOA neurons project to diverse regions, which may have distinct effects on body temperature and blood glucose. (**A**) (*Left*) Schematic for administration of AAV helper virus and HSV anterograde tracer to the avPOA of *Vglut2-IRES-Cre* animals. Mice were first transduced with a complementary helper AAV (AAV-DIO-TK-2A*-*EGFP), and after a two-week recovery period were transduced in the same region with HSV-H129-ΔTK-tdTomato, and sacrificed 65 hours following injection (n=3 *Vglut2-IRES-Cre* animals). (*Right*) (*Top*) Representative image of the starter region of the avPOA and (*Bottom*) one of the downstream targets of avPOA^Vglut2^ neurons in the dorsomedial hypothalamus (DMH). HSV^+^ cells labeled in red, AAV-DIO-TK-EGFP^+^ starter cells labeled in green. Note: in the DMH, HSV^+^ cells predominantly are observed in the ventral portion of the DMH (DMV). (**B)** Representative images of the same regions as Figure 6a in mice transduced with the anterograde monosynaptic tracer HSV-H129-ΔTK-tdTomato. (**C**) Quantitation of HSV expression in (B). Regions were defined in accordance with the Paxinos Reference Atlas, and the number of HSV-positive cells as a fraction of total DAPI cells was calculated for each region (n = 3 *Vglut2-IRES-Cre* animals, mean ±SEM). **(D)** Fold change in plasma glucose concentrations of *PACAP-2A-Cre* animals described in Figure 6i injected with either PBS or CNO at t = 0, 15, 30, 60, and 120 minutes post-injection relative to baseline (** = p < 0.005, student’s two tailed t-test, n = 8 PBS, n = 8 CNO, mean ±SEM). **(E)** Quantification of glucose levels of individual animals from (D) at t = 60 minutes post-injection (** = p < 0.005, student’s two tailed t-test, n = 8 PBS, n = 8 CNO, mean ±SEM).

**Supplementary Figure 6:**
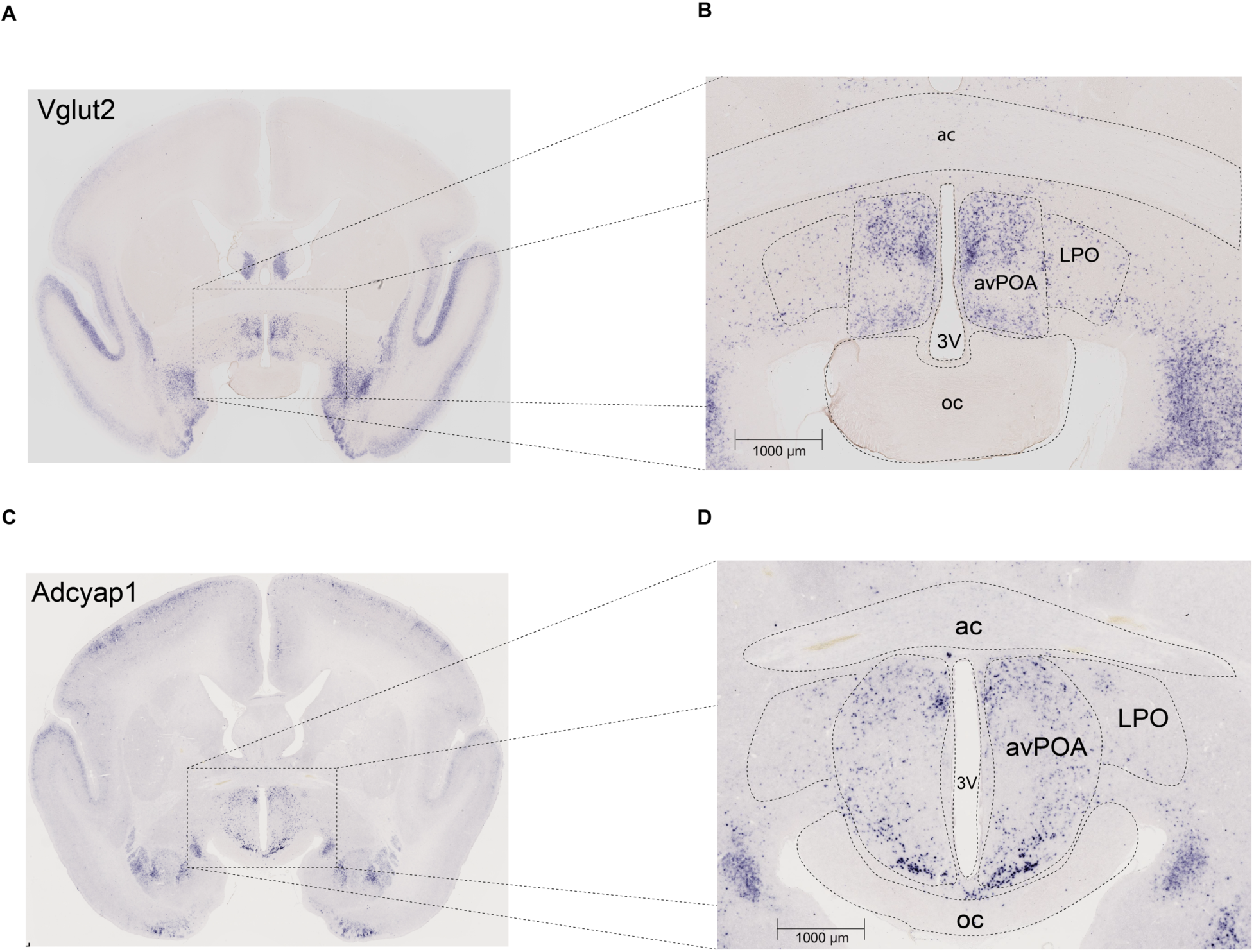
avPOA expression of *Vglut2* and *Adcyap1* in the marmoset brain. **(A)** RNA fluorescence *in situ* hybridization (FISH) image of *Vglut2* RNA expression in coronal marmoset brain region corresponding to mouse avPOA. Purple puncta indicate *Vglut2* RNA expression. **(B)** Magnification of (A) to show the avPOA and LPO regions that correspond to the mouse brain, demonstrating that avPOA^Vglut2^ expression is anatomically conserved between species. **(C)** RNA FISH image of *Adcyap1* RNA expression in coronal marmoset brain region corresponding to mouse avPOA. Purple puncta indicate *Adcyap1* RNA expression. **(D)** Magnification of (C) to show the avPOA and LPO regions that correspond to the mouse brain, demonstrating that avPOA^Adcyap1^ expression is anatomically conserved between species. Images adapted from: https://gene-atlas.brainminds.jp/

